# Synthesis of 4-methylvaleric acid, a precursor of pogostone, involves a 2-isobutylmalate synthase related to 2-isopropylmalate synthase of leucine biosynthesis

**DOI:** 10.1101/2021.09.28.462264

**Authors:** Chu Wang, Ying Wang, Jing Chen, Lang Liu, Zhengguo Li, Eran Pichersky, Haiyang Xu

## Abstract

We show here that the side chain of pogostone, one of the major components of patchouli oil obtained from *Pogostemon cablin* and possessing a variety of pharmacological activities, is derived from 4-methylvaleric acid.
We also show that 4-methylvaleric acid is produced through the one-carbon α- ketoacid elongation pathway with the involvement of the key enzyme 2- isobutylmalate synthase (IBMS), a newly identified enzyme related to isopropylmalate synthase (IPMS) of Leu biosynthesis.
Site-directed mutagenesis identified Met^132^ in the N-terminal catalytic region as affecting the substrate specificity of PcIBMS1. And even though PcIBMS1 possesses the C-terminal domain that in IPMS serves to mediate Leu inhibition, it is insensitive to Leu.
The observation of the evolution of IBMS from IPMS, as well as previously reported examples of IPMS-related genes involved in making glucosinolates in Brassicaceae, acylsugars in Solanaceae, and flavor compounds in apple, indicate that IPMS genes represent an important pool for the independent evolution of genes for specialized metabolism.

**One Sentence Summary:** We describe a novel enzyme, 2-isobutylmalate synthase, that is related to 2-isopropylmalate synthase and is able to efficiently convert 4-methyl-2- oxovalerate to 2-isobutylmalate, a key intermediate in the synthesis of the pogostone precursor 4-methylvaleric acid.

## INTRODUCTION

*Pogostemon cablin* is a traditional Chinese medical plant in the Lamiaceae family, from whose aerial parts *Pogostemonis Herba*, a herbal preparation, is obtained. *Pogostemonis Herba* or its essential oil component (known as patchouli oil) have been widely used in many Asian countries for the treatment of many ailments since ancient time due to a variety of pharmacological activities (Li *et al*., 2013; Wang *et al*., 2016; Hu *et al*., 2017; Chen, JR *et al*., 2021). Pogostone, one of the major chemical components of patchouli oil, has recently been demonstrated to possess various bioactive activities including antimicrobial activities as well as pharmacological activities (Li *et al*., 2012; Huang *et al*., 2014; Li *et al*., 2014; Peng *et al*., 2014; Chen *et al*., 2015; Cao *et al*., 2017; Luchesi *et al*., 2020; Ye *et al*., 2021).

The biosynthetic pathway of pogostone is still unclear, but an outline of the pathway has been proposed that involves the branched-chain 4-methylvaleric acid (Chen, J *et al*., 2021; Fig. 1). Furthermore, PcAAE2, a cytosolic acyl-activating enzyme that catalyzes the conversion of 4-methylvaleric acid to 4-methylvaleryl-CoA as part of the proposed pogostone biosynthetic pathway, has been identified and characterized (Chen, J *et al*., 2021).

**Fig. 1.**
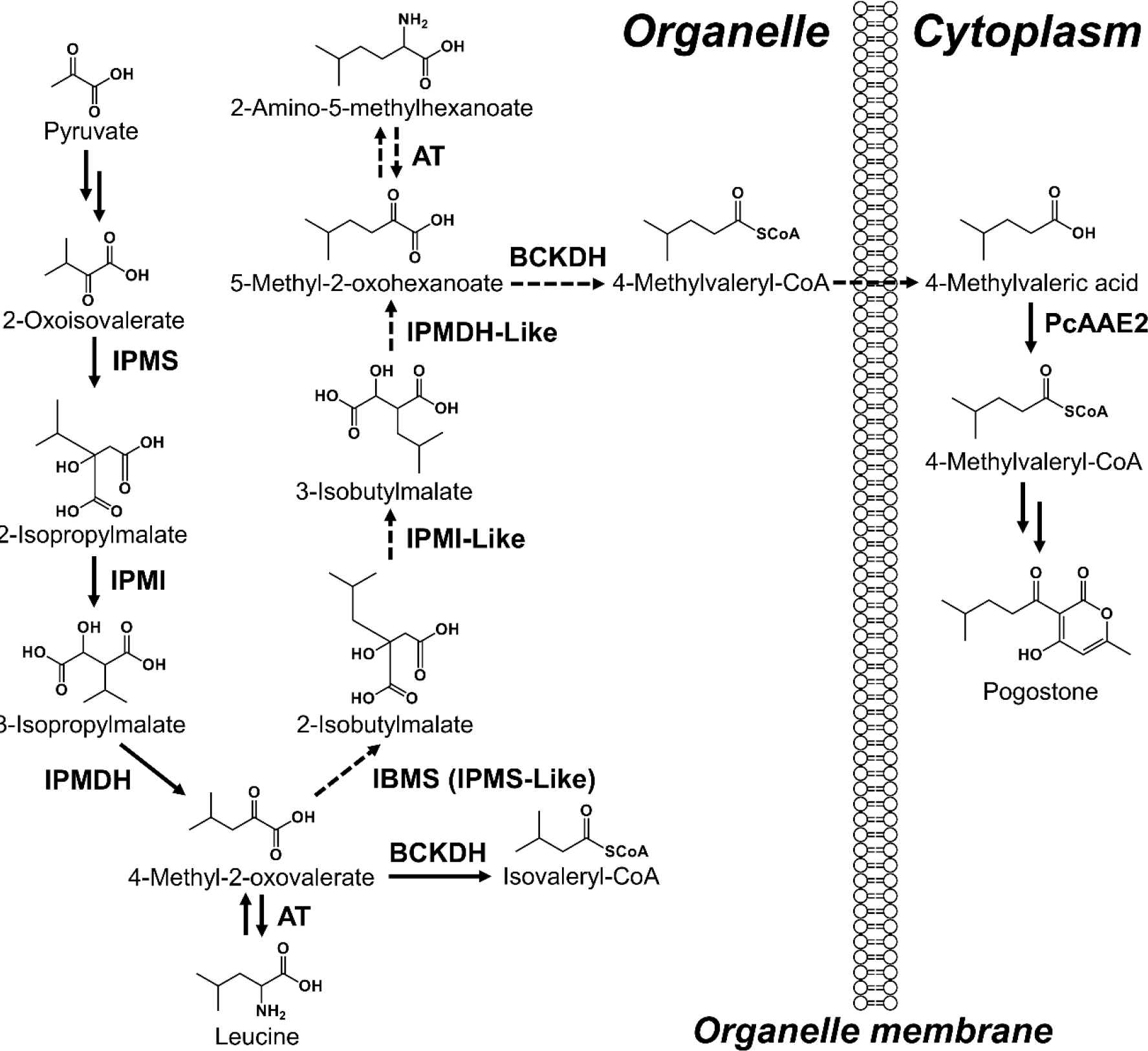
The proposed biosynthetic pathway of 4-methylvaleric acid for pogostone biosynthesis in *Pogostemon Cablin*. The anabolism and catabolism of branched chain amino acid leucine are well documented and occur in plastid and mitochondria, respectively, in land plants (Binder *et al*., 2007; Binder, 2010). 5-Methyl-2-oxohexanoate is elongated by one carbon atom from 4-Methyl-2-oxovalerate which is derived from either anabolism or catabolism of leucine through an α-keto acid elongation (αKAE) pathway. The one-carbon elongation process for 5-Methyl-2-oxohexanoate through αKAE route is performed in three sequential chemical reactions: 2-oxo acid substrate 4-methyl-2- oxovalerate first undergoes an aldol-type condensation with acetyl-CoA by IBMS to give 2-isobutylmalate, which then undergoes isomerization and oxidative decarboxylation by IPMI and IPMDH or their homologies to yield 5-Methyl-2-oxohexanoate. 4-methylvaleryl- CoA is formed from 5-Methyl-2-oxohexanoate in a reaction catalyzed by BCKDH in mitochondria, and then transported into cytosol by a transporter along with losing the CoA group to give free 4-methylvaleric acid, which is subsequently activated again to CoA derivative by the cytosol acyl-activating enzyme 2 (PcAAE2) for pogostone biosynthesis (Chen, J *et al*., 2021). IPMS, 2-Isopropylmalate synthase; IBMS, 2-Isobutylmalate synthase; IPMI, Isopropylmalate isomerase; IPMDH, 3-Isopropylmalate dehydrogenase; AT, Aminotransferase; BCKDH, Branched-chain keto acid dehydrogenase; PcAAE2, Acyl-activating enzyme 2 from *Pogostemon cablin*.

The formation of the branched C6 fatty 4-methylvaleric acid has not yet been elucidated in any species, although evidence for the involvement of either α-ketoacid elongation (αKAE) pathway or the fatty acid synthase (FAS)-mediated elongation pathway in the elongation of other branched-chain fatty acids in various species has been presented (Ohlrogge & Browse, 1995; Kroumova & Wagner, 2003; Slocombe *et al*., 2008; Li-Beisson *et al*., 2013). In the FAS route, both carbons from acetyl-ACP are retained per elongation cycle. In the αKAE pathway, acetyl-CoA is added but only one carbon is retained per elongation cycle. The αKAE pathway plays a role in the biosynthesis of a number of primary and specialized metabolites such as leucine and glucosinolates (Melendez-Hevia *et al*., 1996; de Quiros *et al*., 2000; Halkier & Gershenzon, 2006; Binder, 2010). The αKAE steps involved in leucine biosynthesis have been well documented in various biological organisms (Binder, 2010) and are illustrative of the αKAE route in general (Fig. 1).

In the Solanaceae, many species produced specialized metabolites called acyl sugars, in which at least some of the acyls attached to the sugar moiety are elongated branched- chains (Kroumova & Wagner, 2003; Slocombe *et al*., 2008; Ning *et al*., 2015). Feeding studies with labeled amino acids showed that the branched short chain fatty acid (2- methylpropanoic acid, 3-methylbutanoic acid and 2-methylbutanoic acid) are directly derived from Val, Leu and Ile, respectively (Kroumova & Wagner, 2003), while the branched acyl chains of C6 to C12 are derived from these BCAAs by elongation through α-ketoacid elongation (αKAE) pathway in tobaccos and petunia, but through fatty acid synthase (FAS)-mediated acyl chain elongation pathway in *L. pennellii* and *D. metel* (Kroumova & Wagner, 2003).

Based on its specific length, 4-methylvaleric acid, the branched C6 fatty acid with one more carbon than the leucine intermediate 3-methylbutanoic, is most likely derived from leucine metabolism by one additional elongation cycle of αKAE involving three reactions catalyzed by IPMS, IPMI, and IPMDH, or analogous enzymes (Fig. 1). We show here that the first reaction, in which 4-methyl-2-oxovalerate is elongated to 2-isobutylmalate, is catalyzed by a novel enzyme, 2-isobutylmalate synthase (IBMS), that is closely related to IPMS.

## MATERIALS AND METHODS

### Plant materials and chemicals

The plants of *Pogostemon cablin* and *Nicotiana benthamiana* were grown in soil under the condition as described previously (Chen, J *et al*., 2021). The harvested tissues were flash frozen in liquid nitrogen and stored at -80°C until use.

Since standard 2-isobutylmalate is not available commercially, the substituted 2- malate derivative generated by the condensation of acetyl-CoA to 4-methyl-2-oxovalerate by PcIBMS1 was identified as 2-isobutylmalate based on the exact mass and fragment pattern by LC-QTOF-MS. All other commercial chemicals were purchased from Sigma-Aldrich.

### Isotope feeding study with deuterium-labeled [^2^H11] 4-methylvaleric acid (4MVA- d11)

The five-week-old top leaves (5W-TLeaf) of *P. cablin* plants were placed in a 10-ml feeding solution right before the onset of darkness and transferred to a vacuum desiccator equipped with a vacuum pump. After incubation for 4 mins at the atmospheric pressure of 0.04 Mpa, the leaves in the feeding solution were transferred to a growth chamber and incubated under darkness for 12 h at 25 °C. Both 4-methylvaleric acid (4MVA) and 4MVA- d11 feeding were performed at a concentration of 1 mM. The leaves were cut into pieces and volatiles of 100 mg leaf pieces were extracted with 500 μl MTBE containing 0.01 ng/μl internal standard tetradecane and analyzed by GC-MS. The relative abundances of pogostone and deuterium label incorporated pogostone (pogostone-d11) were calculated by peak area normalization to internal standard tetradecane. 100 μl MTBE extracts were further dried and dissolved in 400 μl 90% methanol for LC-QTOF-MS analysis.

Mass-to-charge ratio (*m/z*) of 224.1 and 235.2 were extracted for pogostone and pogostone-d11, respectively, in GC-MS analysis, while *m/z* of 223.0976 and 234.1666 in negative mode in LC-QTOF-MS analysis were extracted for pogostone and pogostone-d11, respectively. The identification of pogostone-d11 were based on the exact mass and fragment pattern by GC-MS analysis and LC-QTOF-MS analysis.

### Gene identification and coexpression analysis

Identification of gene candidates involved in 4-methylvaleric acid biosynthesis from our previously constructed *P. cablin* transcriptome database (bioproject accession number, PRJNA713906) was performed by querying our local nucleotide database with Arabidopsis representative enzymes involved in BCAA Leu metabolism using the TBLASTN function as previously described (Chen, J *et al*., 2021). Coexpression analysis was performed by comparing the transcript abundance of *PcAAE2* and the identified gene candidates involved in 4-methylvaleric acid biosynthesis using the Pearson correlation method to quantitate the similarity of their expression profiles as described previously (Li *et al*., 2018).

### Quantitative RT-PCR analysis of the identified gene candidates involved in 4- methylvaleric acid biosynthesis

For Quantitative real-time PCR (RT-qPCR) analysis of transcripts in different tissues, RNA was extracted using Total RNA Isolation Kit from Omega with a DNA digestion step. The reverse transcription reaction was performed using High-Capacity cDNA Reverse Transcription Kit from Thermo Fisher Scientific following the manufacturer’s instructions using random primers. The RT-qPCR analysis was performed using Power SYBR Green PCR master mix as previously described (Chen, J *et al*., 2021). The relative expressions of these genes relative to the three reference genes (*GAPDH*, *Actin7* and *Tubulin3*) in each tissue were calculated as previously described (Chen, J *et al*., 2021). Primers used in this study are listed in the Table S1.

**Table 1.**
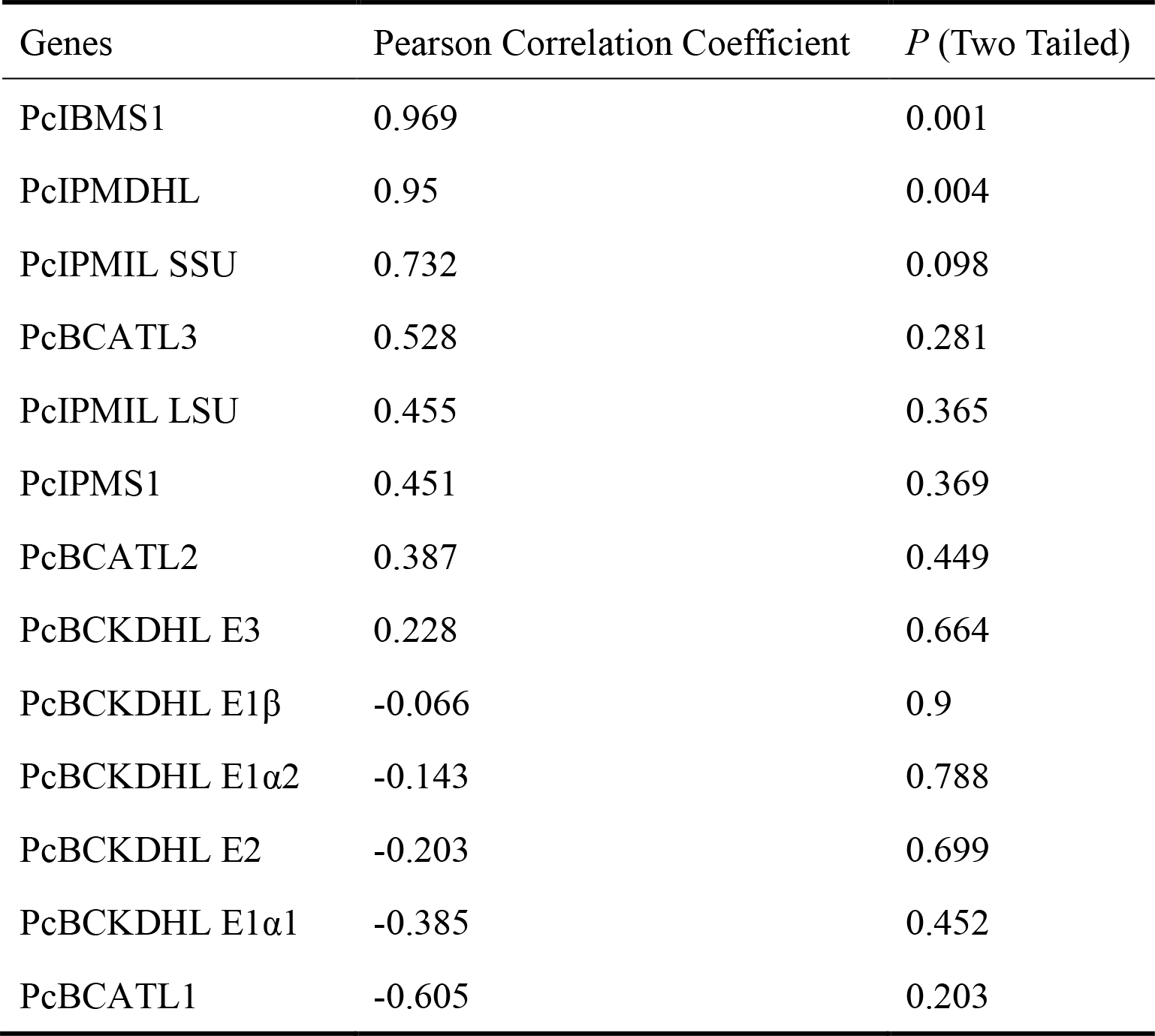
Ranking of 4-methylvaleric acid biosynthetic pathway gene candidates by coexpression correlation analysis with the *PcAAE2* gene.

### Isolation of His-tagged recombinant proteins of PcIBMS1, PcIPMS1, PcIBMS1 M132L and PcIPMS1 L135M

To obtain soluble proteins for expression in *Escherichia coli*, the truncated open reading frames of *PcIBMS1* and *PcIPMS1*, both missing the first 40 amino acids, were obtained by Reverse transcriptase polymerase chain reaction (RT-PCR) from prepared cDNA of *P. cablin* seedlings and were introduced into the expression vector pET-28a (+) between the restriction sites of NdeI and BamHI using NEBuilder® HiFi DNA Assembly Cloning Kit (NEB, Catalog number: E5520S), in each case generating a fusion gene that encoded a “tag” of HIS6 residues at the N-terminus for expression in *E. coli*. The corresponding resulted expression vectors are pET-28a-PcIBMS1 and pET-28a-PcIPMS1, respectively. The two fragments for assembly for PcIBMS1 M132L and PcIPMS1 L135M constructs were obtained by RT-PCR from corresponding vectors pET-28a-PcIBMS1 and pET-28a-PcIPMS1 and were assembled into the expression vector pET-28a (+) in the same way as the construction of pET-28a-PcIBMS1 and pET-28a-PcIPMS1. The corresponding resulted expression vectors are pET-28a-PcIBMS1 M132L and pET-28a-PcIPMS1 L135M, respectively. Recombinant proteins of PcIBMS1, PcIPMS1, PcIBMS1 M132L and PcIPMS1 L135M were expressed in *E. coli* strain BL21(BE3) and purified using Ni-NTA agarose chromatography (Qiagen) as previously described (Xu et al., 2018). The primer pairs used for construction of these pET-28a (+) expression vectors fused with HIS6 residues at the N-terminus for expression in *E. coli* are listed in Table S1.

### Enzymatic assays of recombinant PcIBMS1, PcIPMS1, PcIBMS1 M132L and PcIPMS1 L135M

The enzyme assay for the condensation reaction between acetyl-CoA and different 2- oxo acids was performed as previously described (de Kraker *et al*., 2007) with minor modification. Briefly, substrate specificity assays for recombinant PcIBMS1 and PcIPMS1 were determined with a spectrophotometric end-point assay with DTNB (Sigma-Aldrich). An aliquot of 1-5 μL of the protein preparation was added to an enzyme assay mixture (100 mM Tris, pH 8.0, 1 mM acetyl-CoA, 10 mM MgCl2 and 0.3 mM 2-oxo acid) for a final volume of 200 μL and incubated for 10 min at 25 °C. The reaction was stopped by freezing in liquid nitrogen and subsequent addition of 160 μL ethanol and 80 μL of 2 mM DTNB (fresh solution in 100 mM Tris, pH 8.0). The mixture was left for 2 to 3 min for DTNB to react with free thiol groups released from acetyl-CoA to create a yellow-color TNB^2-^ dianion product with an ε412 of 14140 M^-1^ cm^-1^ (Kohlhaw, 1988). After full color development, the absorbance of the mixture was measured at 412 nm and then adjusted by subtracting the background of the identical enzyme assay mixture without 2-oxo acid. For 4-Methylthio-2-oxobutanoic acid, the results were further corrected for the slight reactivity between its thiol group and DTNB using reactions containing the 2-oxo acid, but not acetyl- CoA. The assay was generally linear for the first 15 min with 1 μg of protein.

Steady-state kinetic assays were DTNB end-point assays that run at 25 °C using 1 to 5 μL aliquots of protein preparation. Absorbance of the reaction was found to increase linearly for at least 15 min. For individual assays, absorbance was recorded every 5 min for 10 min. Correction for background absorbance was performed as described above. Each assay was repeated 3 times. Kinetic assays for 2-oxo acid were performed with concentration of acetyl-CoA fixed at 1 mM while the concentration of 2-oxo acid ranging from 0.05 to 4 mM. Kinetic assays for acetyl-CoA were performed with the concentration of 2-oxo acid fixed at 2 mM while the concentration of acetyl-CoA ranging from 0.02 mM to 1 mM. Kinetic parameters were determined using the hyperbolic regression analysis method in Hyper32 software (version 1.0.0, http://hyper32.software.informer.com/). The spectrophotometric assay was also used to test the effect of Leu on enzyme activity in the range from 0.05 mM to 10 mM.

In addition, the condensation activities of recombinant PcIBMS1 and PcIPMS1 towards 2-oxoisovalerate and 4-methyl-2-oxovalerate were further measured using an LC- QTOF-MS-based method. Briefly, the reaction mixtures (200 μL) contain 0.1 mM Tris- HCl, pH 8.0, 0.5 mM acetyl-CoA, 10 mM MgCL2, 0.5 mM 2-oxo acid substrate. The reaction was initiated by adding 2 μg purified enzyme protein and stopped after 20 min by addition of 400 μL methanol. The solution was filtered through a 0.45 μm filter prior to LC-QTOF-MS analysis.

### Subcellular Localization of PcIBMS1 and PcIPMS1

The vector used for subcellular localization of analysis was pTF486. The construction of these vectors expressing fusion proteins of PcIBMS1 and PcIPMS1 with GFP fused at C-terminus was performed as previously described (Chen, J *et al*., 2021). The full-length open reading frames of PcIBMS1 and PcIPMS1 were cloned into pTF486 between SalI and BamHI sites using In-Fusion® HD Cloning Kit (TaKaRa) following the protocol. Protoplasts were prepared from Arabidopsis leaves, and transformation and confocal microscopy were performed as described previously (Yoo *et al*., 2007). Primers used in the construction of these vectors are listed in Table S1.

### Transient expression in *N. benthamiana* leaves

The pEAQ-HT constructs carrying each of *PcIBMS1* and *PcIPMS1* used for transformation were constructed as described previously (Xu *et al*., 2019). The procedure used for transient expression in *N. benthamiana* leaves was performed as previously described (Xu *et al*., 2019). Leaves of 4-week-old *N. benthamiana* plants were used for infiltration. *N. benthamiana* plants transformed with empty pEAQ-HT vector were used as control. The infiltrated leaves were collected 7 days after infiltration.

To analyze the compounds produced in leaves, 500 mg of ground leaf tissues were mixed thoroughly with 100 µl ultrapure water and then centrifuged to remove pellet. 50 µl supernatant were mixed thoroughly with 450 µl methanol containing 0.01mM naringenin and filtered through a 0.45 μm filter for LC-QTOF-MS analysis. 200 µl supernatant were extracted with equal volume of MTBE containing 10 ng/µl geraniol as internal standard. The MTBE extracts were then analyzed by GC-MS.

Base hydrolysis of modified branched chain fatty acids was performed as previously described (Xu *et al*., 2018). 500 mg of ground leaf tissues were mixed completely with 50 µL 4N NaOH and incubated at 80°C for 20 min, followed by neutralization with 50 µL 4N HCl, and then centrifuged at 10000 g for 15 mins. 200 µl supernatant were extracted with equal volume of MTBE containing 10 ng/µl geraniol for GC-MS analysis.

### GC-MS analysis and LC-QTOF-MS analysis

For GC-MS analysis, 1 µl aliquot of the sample was injected into Agilent GC-MSD (Agilent 7890B-5977B) system equipped with the Rxi-5Sil MS column (30 m × 0.25 mm × 0.25 μm film thickness, RESTEK, USA). The oven temperature was programmed as previously described (Chen, J *et al*., 2021). For LC-QTOF-MS analysis, 2 µl aliquot of the sample was injected into LC-QTOF-MS system (Agilent, 6545 LC/QTOF-MS) coupled with a C18 column (ZORBAX RRHD Plus C18, Φ2.1 × 50mm, 1.8 μm) at 35°C. The gradient (solvent A, water + 0.1% formic acid; solvent B, acetonitrile) program and operating parameters were set as previously described (Chen, J *et al*., 2021).

## RESULTS

### 4-Methylvaleric acid is a precursor of pogostone

Chen, J et al (2021) hypothesized that the side chain of pogostone is derived from 4- methylvaleric acid, and furthered showed that the expression of PcAAE2, a cytosolic acyl-activating enzyme that catalyzes the conversion of 4-methylvaleric acid to 4- methylvaleryl-CoA, is correlated with pogostone biosynthesis. However, a direct proof that 4-methylvaleric acid is a precursor of pogostone has not yet been presented. To determine if the side chain of pogostone is derived from 4-methylvaleric acid, five-week-old top leaves (5W-TLeaf) of *P. cablin* plants were placed in a solution containing the deuterium-labeled [^2^H11] 4-methylvaleric acid (4-methylvaleric-d11 acid or 4MVA-d11) at the concentration of 1 mM (Fig. 2). Following an overnight treatment with the labeled 4MVA- d11, volatiles were extracted with MTBE and aliquots were analyzed by gas chromatography-mass spectrometry (GC-MS). Another portion of each MTBE extract was further dried and dissolved in 90% methanol for liquid chromatography-quadrupole time-of-flight mass spectrometry (LC-QTOF-MS). We observed that 7.8% of the pogostone molecules extracted had an 11-D shift (from 223.0976 for pogostone to 234.1666 for pogostone-d11 in negative mode in LC-QTOF-MS analysis and from 224.1 for pogostone to 235.2 for pogostone-d11 in GC-MS analysis) in the parent ion mass (Fig. 2), indicating that 4-methylvaleric acid is a bona fide precursor of pogostone biosynthesis.

**Fig. 2.**
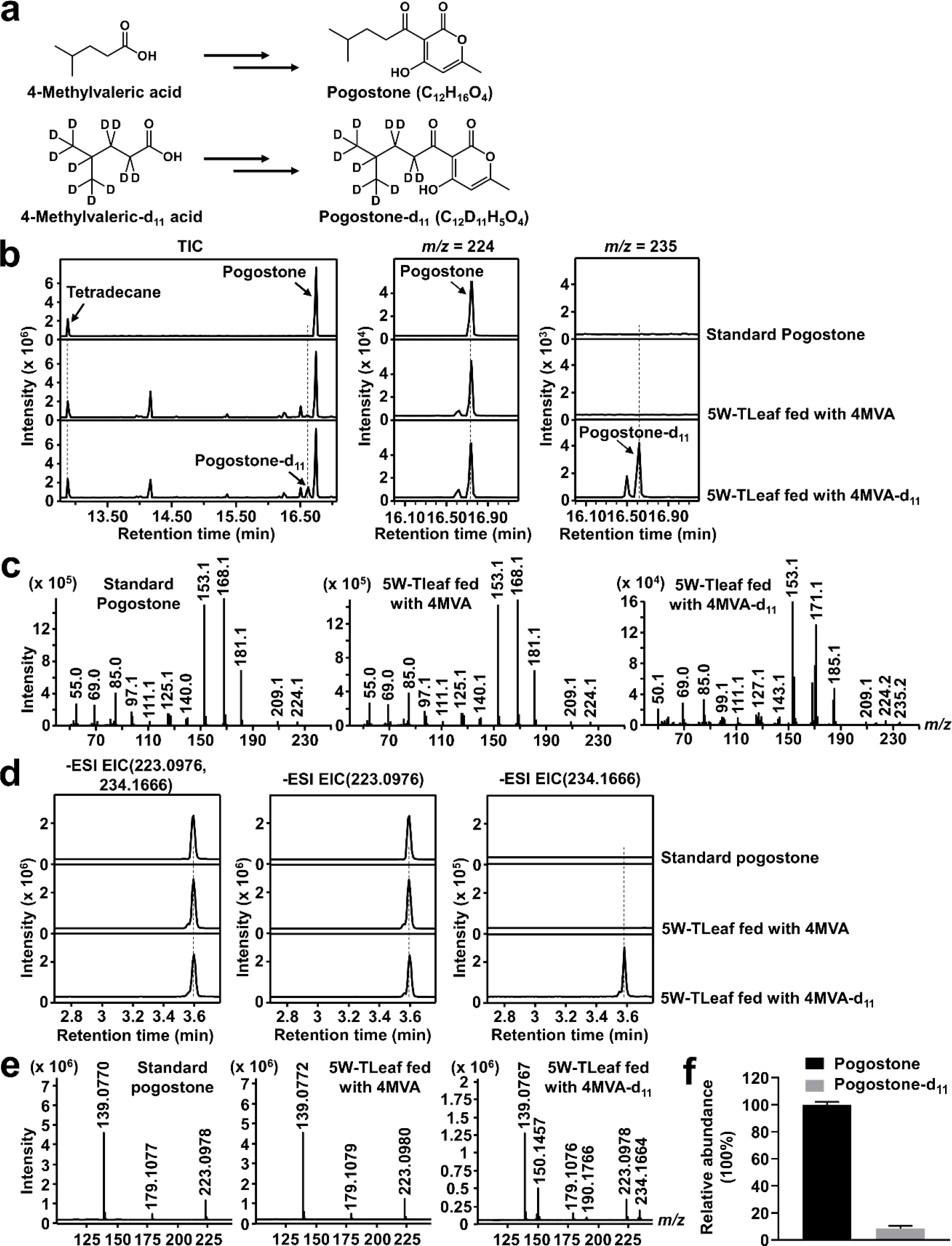
Stable isotope incorporation into pogostone from *Pogostemon cablin* leaves fed with 4-methylvaleric-d11 acid. **a.** The chemical structures of pogostone from 4- methylvaleric acid (4MVA) and 4-methylvaleric-d11 acid (4MVA-d11). **b.** GC-MS analysis of *P. cablin* five-week-old top leaf (5W-TLeaf) fed with 4MVA or 4MVA-d11. The total ion chromatograms (TIC) (left) and extracted ion chromatograms of *m/z* 224 for pogostone (middle) and of *m/z* 235 for pogostone-d11 (right) are shown. Tetradecane was used as internal standard. **b.** The mass spectrums in GC-MS analysis of standard pogostone (upper left), pogostone from *P*. *cablin* 5W-TLeaf fed with 4MVA (upper right) and pogostone-d11 from *P*. *cablin* 5W-TLeaf fed with 4MVA-d11 (lower left). Note the 11-mass-unit shift of the parent ions of pogostone (224) to pogostone-d11 (235) upon feeding with 4MVA-d11. **d.** LC-QTOF-MS analysis of *P. cablin* five-week-old top leaf (5W-TLeaf) fed with 4MVA or 4MVA-d11. The extracted ion chromatograms in negative mode of *m/z* 223.0976 and 234.1666 (left) for both pogostone and pogostone-d11, 223.0976 (middle) for pogostone and 234.1666 (right) for pogostone-d11 are shown. **e.** The mass spectrums in negative mode in LC-QTOF-MS analysis of standard pogostone (left), pogostone from *P*. *cablin* 5W- TLeaf fed with 4MVA (middle) and pogostone-d11 from *P*. *cablin* 5W-TLeaf fed with 4MVA-d11 (right). Note the 11-mass-unit shift of the parent ions of pogostone (223.0976) to pogostone-d11(234.1666) upon feeding with 4MVA-d11. **f.** The relative abundances of pogostone and pogostone-d11 in *P*. *cablin* 5W-TLeaf fed with 4MVA-d11. The relative abundances in GC-MS analysis of pogostone and pogostone-d11 in *P*. *cablin* 5W-TLeaf fed with 4MVA-d11 were calculated by peak area normalization to internal standard tetradecane and the relative abundance of pogostone was set as 100%. Data are represented as mean ± SD of three independent biological replicates.

### Identification of the gene candidates involved in 4-methylvaleric acid biosynthesis by coexpression analysis

To identify the candidate genes involved in 4-methylvaleric acid biosynthesis, we performed homology searches on our previously constructed *P. cablin* RNAseq database (Chen, J *et al*., 2021), using genes involved in leucine metabolism in *Arabidopsis thaliana* using TBLASTN software (Fig. 1). We obtained a total 13 unique genes (Table S2): two genes with similarity to AtIPMS (named *PcIBMS1* and *PcIPMS1, see below)*; two genes (*PcIPMIL SSU* and *PcIPMIL LSU; “L” for “-like”)* related to the small subunit and large subunit of Arabidopsis IPMI, respectively; one gene (*PcIPMDHL)* related to AtIPMDH; three genes putatively encoding branched-chain aminotransferases (*PcBCATL1*, *PcBCATL2* and *PcBCATL3)*; and five genes putatively encoding the three subunits of branched chain *α-*keto acid dehydrogenase (*PcBCKDHL E1α1*, *PcBCKDHL E1α2*, *PcBCKDHL E1β*, *PcBCKDHL E2* and *PcBCKDHL E3)*.

**Table 2.**
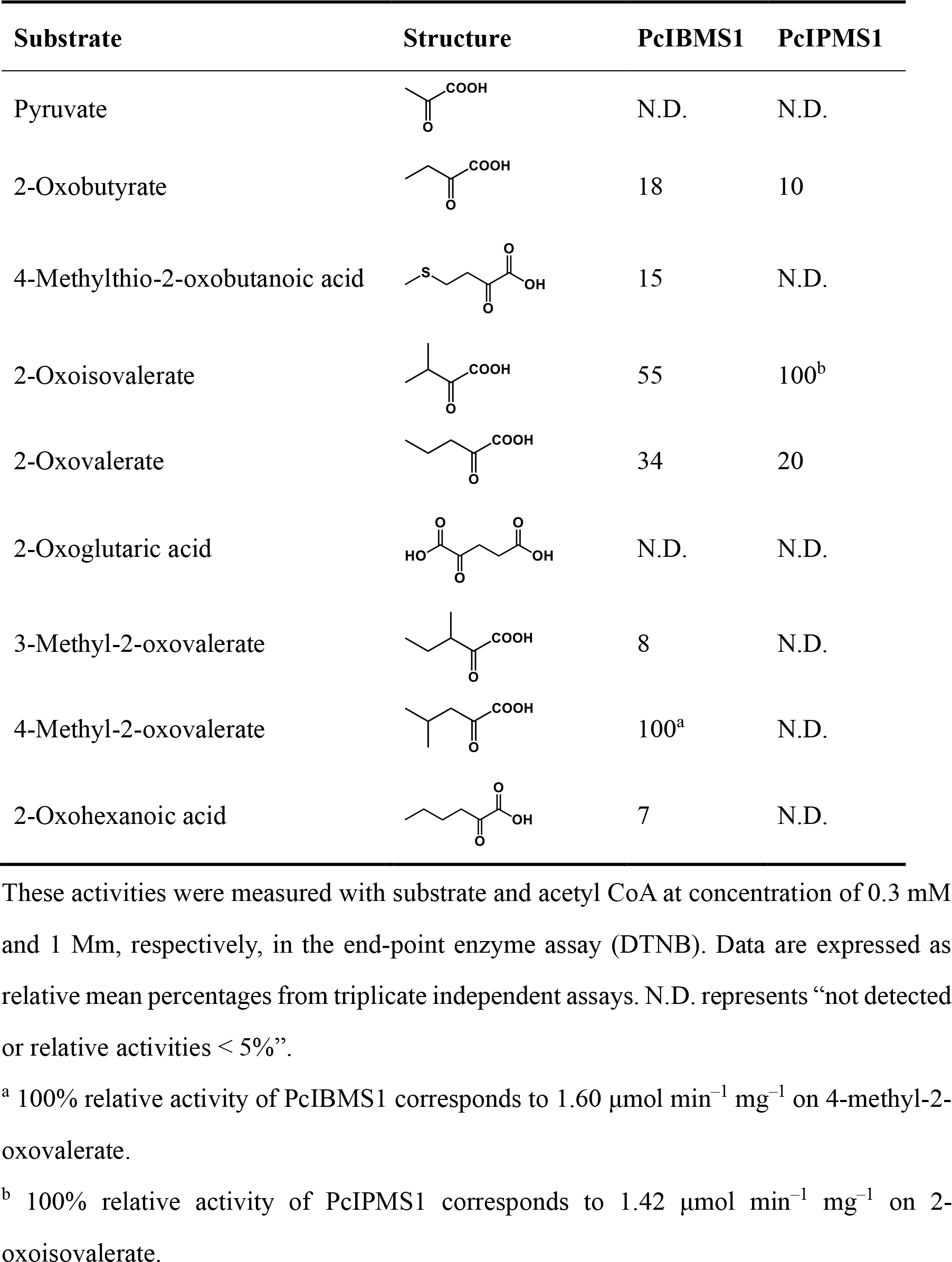
Substrate Specificity of Recombinant PcIBMS1 and PcIPMS1.

| Substrate | Structure | PcIBMS1 | PcIPMS1 |
| --- | --- | --- | --- |
| Pyruvate |  | N.D. | N.D. |
| 2-Oxobutyrate |  | 18 | 10 |
| 4-Methylthio-2-oxobutanoic acid |  | 15 | N.D. |
| 2-Oxoisovalerate |  | 55 | 100<sup>b</sup> |
| 2-Oxovalerate |  | 34 | 20 |
| 2-Oxoglutaric acid |  | N.D. | N.D. |
| 3-Methyl-2-oxovalerate |  | 8 | N.D. |
| 4-Methyl-2-oxovalerate |  | 100<sup>a</sup> | N.D. |
| 2-Oxohexanoic acid |  | 7 | N.D. |

These activities were measured with substrate and acetyl CoA at concentration of 0.3 mM and 1 Mm, respectively, in the end-point enzyme assay (DTNB). Data are expressed as relative mean percentages from triplicate independent assays. N.D. represents “not detected or relative activities < 5%”.
<sup>a</sup> 100% relative activity of PcIBMS1 corresponds to 1.60 $\mu\text{mol min}^{-1} \text{mg}^{-1}$ on 4-methyl-2-oxovalerate.
<sup>b</sup> 100% relative activity of PcIPMS1 corresponds to 1.42 $\mu\text{mol min}^{-1} \text{mg}^{-1}$ on 2-oxoisovalerate.

We next performed coexpression analysis of these 13 unique genes through Pearson correlation analysis to look for genes whose expression was best correlated with that of *PcAAE2*, whose expression was previously shown to be correlated with pogostone biosynthesis (Chen, J *et al*., 2021), in all six *P. cablin* tissues including Seedling, 5W-Root, 5W-Stem, 5W-TLeaf, 8W-TLeaf and 8W-SLeaf (Table S3). Two of the 13 genes, *PcIBMS1* and *PcIPMDHL*, showed statistically significance correlation (*P* < 0.01) (Table 1). In addition, in the coexpression analysis of all the unique genes in the *P. cablin* RNAseq database with the *Pc*AAE2 gene through Pearson correlation analysis, *PcIBMS1* ranked the top third unique gene (Table S4).

**Table 3.**
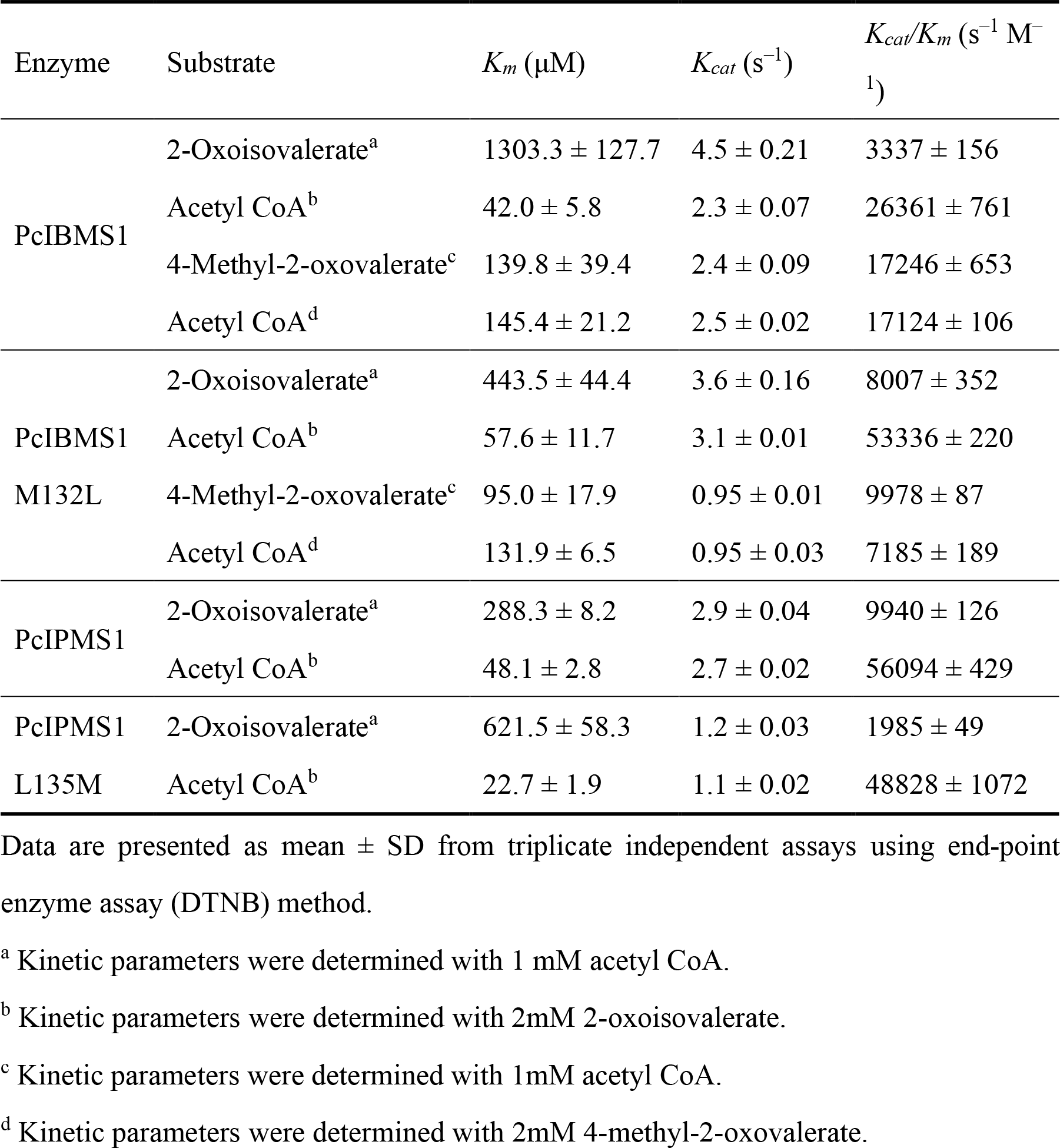
Kinetic Properties of Recombinant PcIBMS1, PcIPMS1, PcIBMS1 M132L and PcIPMS1 L135M.

To more accurately test whether the transcript profiles of these 13 genes correlated with that of *PcAAE2* in different *P. cablin* tissues at different developmental stages as previously described (Chen, J *et al*., 2021), their expression patterns were further confirmed by RT-qPCR in the same 13 tissue samples used for *PcAAE2* (Fig. S1). Again, the expression patterns of *PcIBMS1* in the tested tissue samples showed a strong positive correlation with that of *PcAAE2*, displaying a similar developmental pattern as that of *PcAAE2* and the distribution pattern of pogostone (Chen, J *et al*., 2021).

### Subcellular localization of PcIBMS1 and PcIPMS1

To investigate the subcellular compartmentation of PcIBMS1, subcellular localization prediction software TargetP 2.0 was first utilized for signal peptide prediction of PcIBMS1 as well as for PcIPMS1. Both PcIBMS1 and PcIPMS1 were predicted to harbor chloroplast transit peptides in their N-termini with a high score (Table S2). The subcellular localizations of these two proteins were further analyzed experimentally by transiently expressing in Arabidopsis leaf protoplasts PcIBMS1 and PcIPMS1 each fused to the green fluorescent protein (GFP) at their C-termini. In these experiments, complete co-localization of both PcIBMS1-GFP and PcIPMS1-GFP signals with red chlorophyll autofluorescence was observed (Fig. 3), indicating that both proteins are targeted to the plastids.

**Fig. 3.**
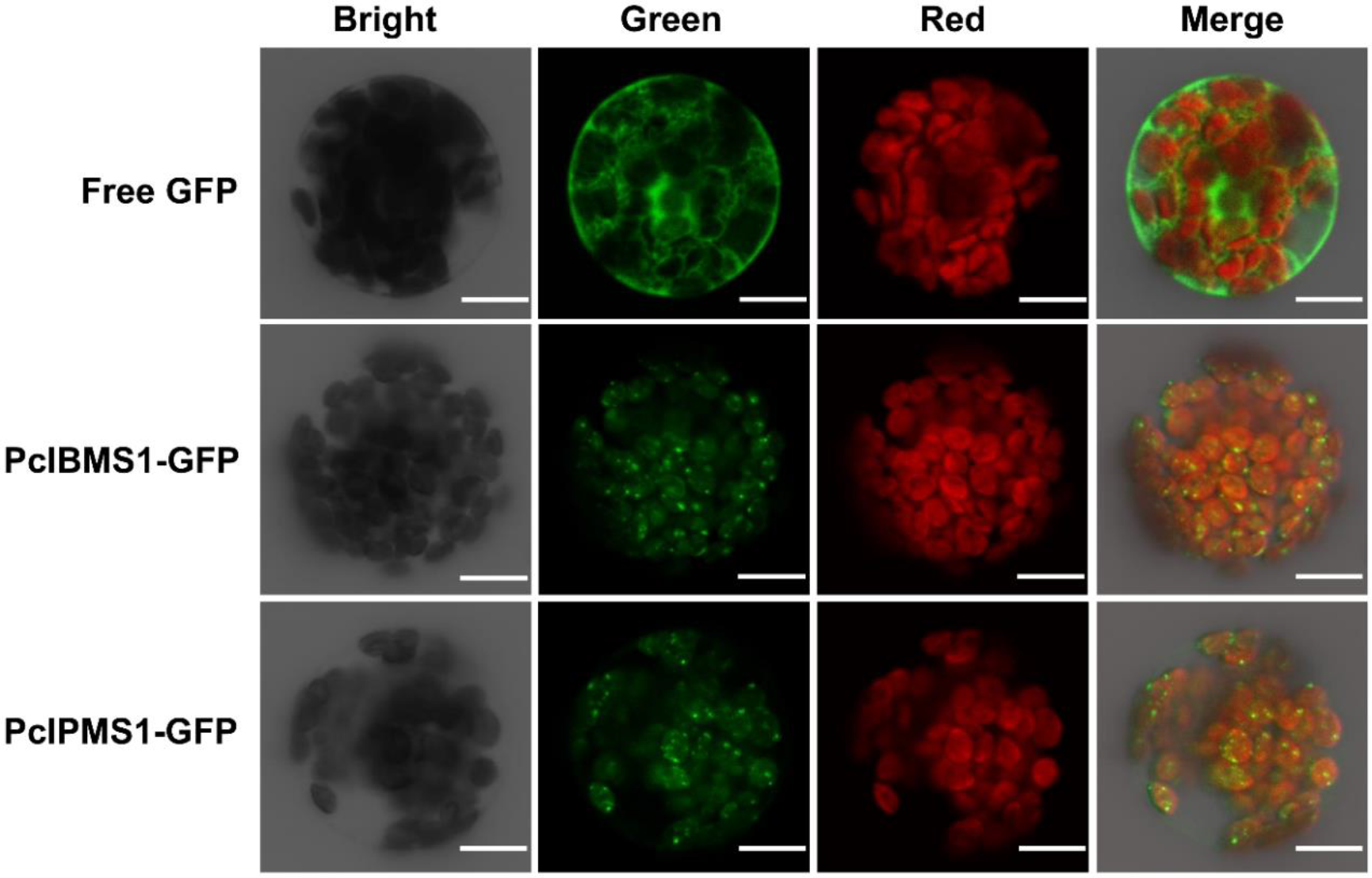
Subcellular localization of PcIBMS1 and PcIPMS1 in Arabidopsis leaf mesophyll protoplasts revealed by laser confocal microcopy. **The signal of free GFP** was used as control and chloroplasts are revealed by red chlorophyll autofluorescence. Scale bar = 10 μm.

### *In vitro* characterization of isopropylmalate and isobutylmalate synthase activities of PcIBMS1

The combination of the positive correlation of expression patterns of *PcIBMS1* with that of *PcAAE2* and its further confirmation by RT-qPCR analysis made it the most likely candidate gene encoding the key enzyme in the biosynthetic pathway of 4-methylvaleric acid compared with the other twelve unique genes. To test this hypothesis, we performed *in vitro* biochemical assays with PcIBMS1 and, for comparison, PcIPMS1. Truncated PcIBMS1 and PcIPMS1 proteins, both lacking the first 40 amino acids (the estimated length of the transit peptide), were produced in *Escherichia coli* with a fused N-terminal His6 tag, and purified as soluble proteins. Both purified recombinant proteins were initially tested in a spectrophotometric end-point enzyme assay (DTNB, 5,5’ -dithiobis (2- nitrobenzoic acid)) with a variety of 2-oxo acid substrates at a 0.3 mM concentration of the 2-oxo acid substrate and 1mM of acetyl-CoA.

Recombinant PcIBMS1 showed strong activity with 4-methyl-2-oxovalerate as a substrate, and also had significant activity with 2-oxoisovalerate (55% of its activity with 4-methyl-2-oxovalerate), but lower or no activity with the other substrates tested (Table 2). Recombinant PcIPMS1 utilized 2-oxoisovalerate as the best substrate, with only low levels of activity or no activity with all other substrates tested. (Table 2). In particular, PcIPMS1showed no detectable activity with 4-methyl-2-oxovalerate (Table 2). The non-availability of commercial larger analogs of 4-methyl-2-oxovalerate such as 5-methyl-2- oxo-hexanoate prevents us from testing enzymatic activities of PcIBMS1 and PcIPMS1 towards them.

To further characterize PcIBMS1 and PcIPMS1, we first tested the dependency of their enzymatic activities on pH and cations. The highest levels of activities of PcIBMS1 and PcIPMS1 with their respective preferred substrates were observed at pH 8.0 (Fig. S2). The activities of PcIBMS1 and PcIPMS1 were dependent on divalent cations with Mg^2+^ being the most preferred cation at optimal concentration of 10 mM (Fig. S2). The kinetic properties of PcIBMS1 with 4-methyl-2-oxovalerate and 2-oxoisovalerate as substrates, and PcIPMS1 with 2-oxoisovalerate as the substrate were measured in a spectrophotometric end-point enzyme assay. The kinetic analysis revealed that PcIBMS1 had a *Km* value of 139.8 ± 39.4 µM and a catalytic efficiency of 1.72 × 10^4^ s^-1^M^-1^ for 4- methyl-2-oxovalerate and a *Km* value of 1303.3 ± 127.7 µM and a catalytic efficiency of 3.337 × 10^3^ s^-1^M^-1^ for 2-oxoisovalerate, and that PcIPMS1 had a *Km* value of 288.3 ± 8.2 µM and a catalytic efficiency of 9.94 × 10^3^ s^-1^M^-1^ for 2-oxoisovalerate (Table 3). The enzymatic products of PcIBMS1 and PcIPMS1 with 2-oxoisovalerate and 4-methyl-2- oxovalerate as substrates were further verified with LC-QTOF-MS (Fig. 4).

**Fig. 4.**
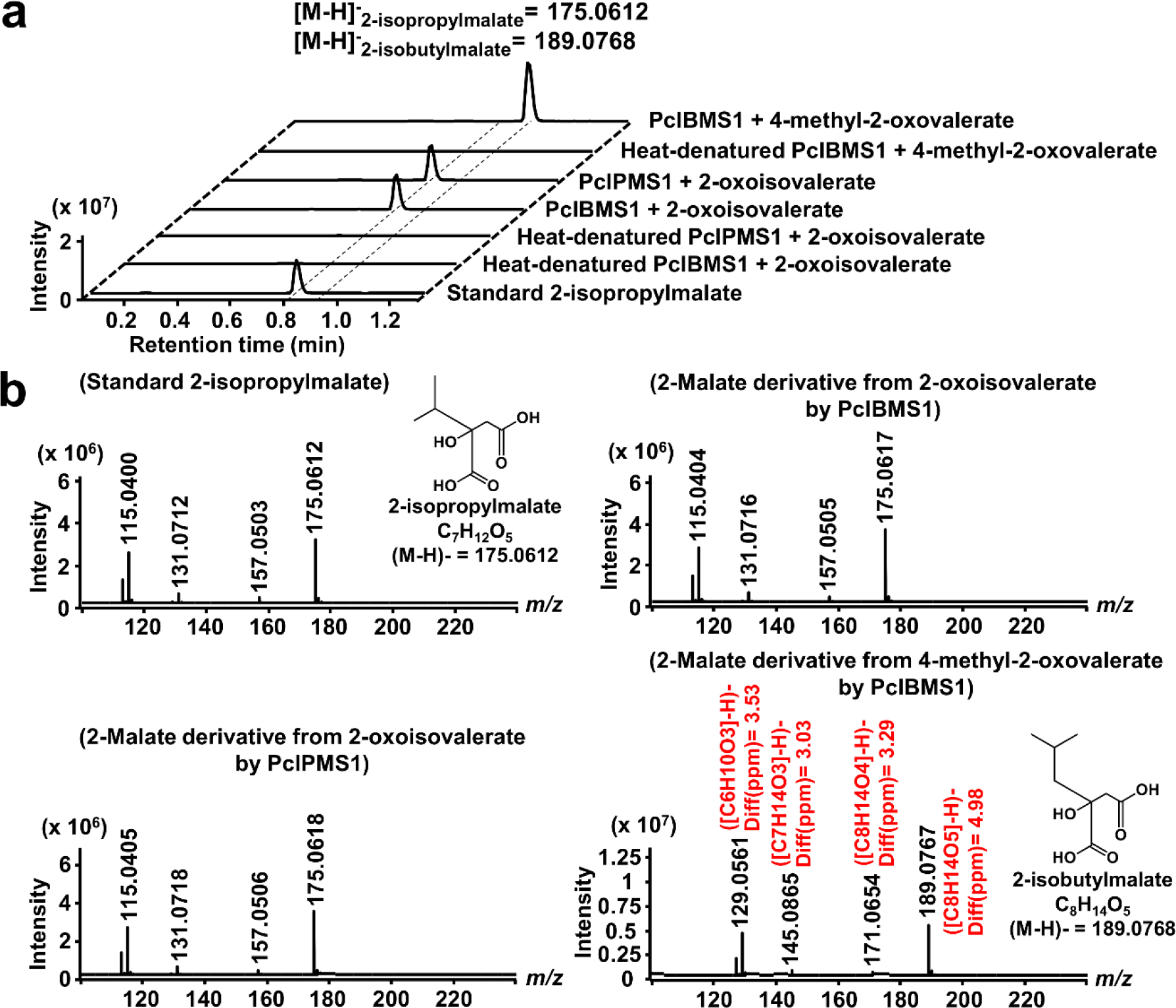
*In vitro* analyses of PcIBMS1 and PcIPMS1 activities. **a.** LC-QTOF-MS analysis of 2-malate derivatives formed in reactions containing purified PcIBMS1 and PcIPMS1, acetyl-CoA, MgCl2, 2-oxoisovalerate or 4-methyl-2-oxovalerate after 30 min of incubation. The extracted ion chromatograms in negative mode of *m/z* 175.0612 and 189.0768 are shown for 2-isopropylmalate and 2-isobutylmalate respectively. **b.** Mass spectrums in negative mode of standard 2-isopropylmalate, and 2-malate derivatives (predicted 2- isopropylmalate and 2-isobutylmalate) generated *in vitro* by the action of PcIBMS1 and PcIPMS1 with the substrates 2-oxoisovalerate or 4-methyl-2-oxovalerate. The 2-malate derivatives from 2-oxoisovalerate by the action of PcIBMS1 and PcIPMS1 were identified as 2-isopropylmalate through the comparison of its retention time and mass spectrums with that of standard 2-isopropylmalate. The 2-malate derivative from 4-methyl-2-oxovalerate by the action of PcIBMS1 was identified as 2-isobutylmalate based on the exact mass and fragment pattern. The chemical structures of 2-isopropylmalate and 2-isobutylmalate were placed along with the mass spectrums of standard 2-isopropylmalate and predicted 2- isobutylmalate respectively. The predicted molecular formula corresponding to the mass fragment of 2-malate derivative generated *in vitro* by the action of PcIBMS1 with the substrate 4-methyl-2-oxovalerate was marked in red color.

### Heterologous expression of PcIBMS1 and PcIPMS1 in Nicotiana benthamiana

To further test the activities of PcIBMS1 and PcIPMS1 *in planta*, *N. benthamiana* leaves were infiltrated with *Agrobacterium tumefaciens* strains harboring plasmids containing either *PcIBMS1* or *PcIPMS1*. For control, *Agrobacterium tumefaciens* strains harboring empty vector (pEAQ-HT) were infiltrated into *N. benthamiana* leaves. In these experiments, the complete open reading frames of *PcIBMS1* and *PcIPMS1* were used, including the corresponding transit peptides that were shown to direct the protein to the plastid. Transformed leaves were harvested seven days after infiltration and the products were extracted and analyzed by LC-QTOF-MS. As expected, transient expression of *PcIBMS1* in *N. benthamiana* leaves resulted in the production of 2-isobutylmalate and 2- isopropylmalate, which were verified by the comparison of their retention times and mass spectrums with corresponding standards (Fig. 5a). In contrast, transient expression of *PcIPMS1* in *N. benthamiana* leaves resulted in negligible amounts of 2-isopropylmalate and 2-isobutylmalate compared with that produced in *N. benthamiana* leaves transiently expressing *PcIBMS1* (Fig. 5a).

**Fig. 5.**
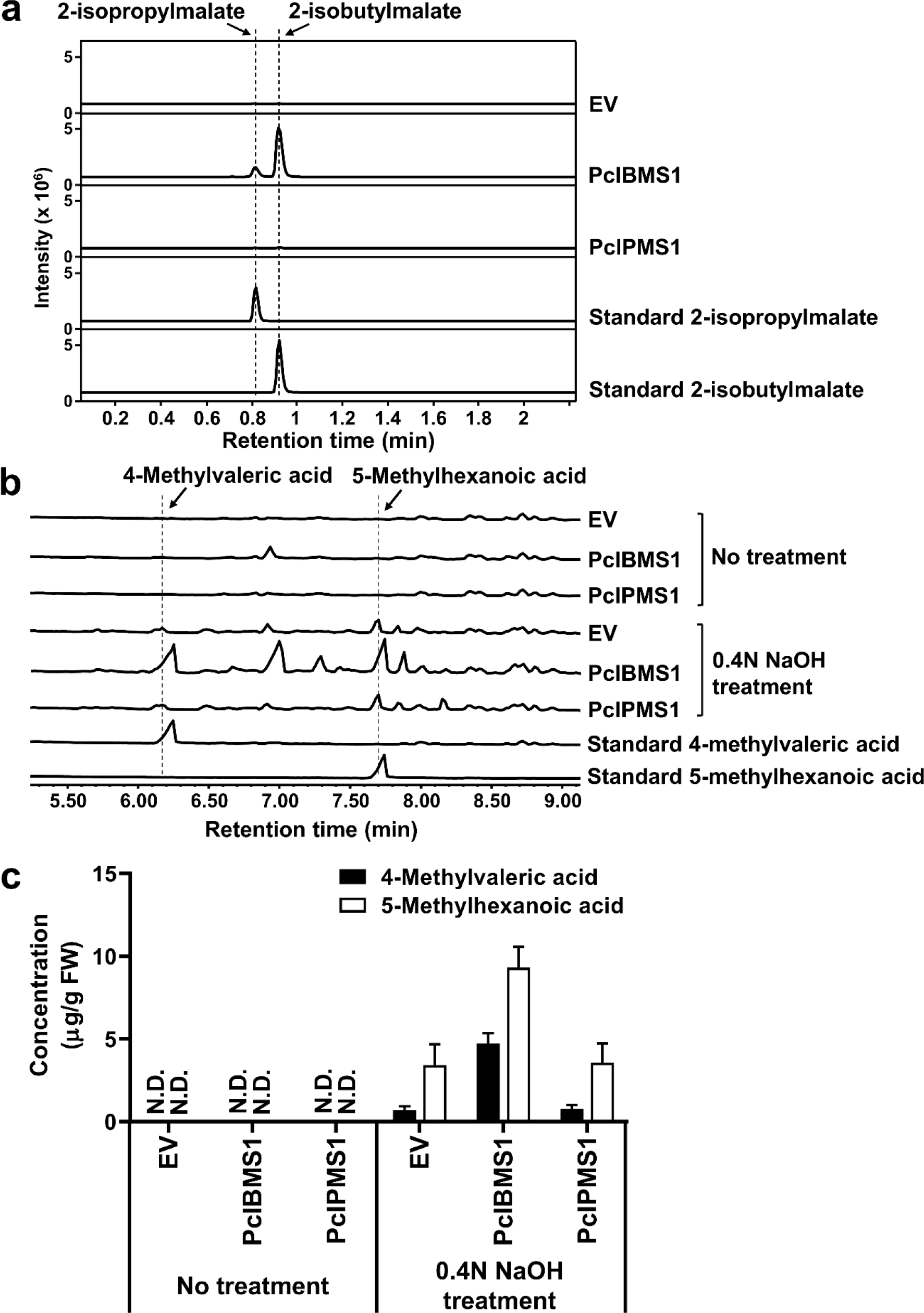
LC-QTOF-MS and GC-MS analyses of *N. benthamiana* leaves expressing *PcIBMS1* and *PcIPMS1*. **a.** LC-QTOF-MS analyses of *N. benthamiana* leaves expressing the two enzymes PcIBMS1 and PcIPMS1. *N. benthamiana* leaves transformed with empty vector (EV) were used as negative control. Extracted ion chromatograms in negative mode of both *m/z* 175.0612 for 2-isopropylmalate and 189.0768 for 2-isobutylmalate are shown. Standard 2-isopropylmalate is available commercially while standard 2-butylmalate was generated by the *in vitro* enzymatic reactions of purified PcIBMS1 with 4-methyl-2- oxovalerate as the substrate in the presence of acetyl-CoA. **b.** GC-MS analyses of *N. benthamiana* leaves expressing the two enzymes PcIBMS1 and PcIPMS1 with or without base treatment. The MTBE extracts of *N. benthamiana* leaf samples with or without base treatment were analyzed by GC-MS and total ion chromatograms are shown. **c.** Concentration of 4-methylvaleric acid and 5-methylhexanoic acid in *N. benthamiana* leaves expressing the two enzymes PcIBMS1 and PcIPMS1 with or without base treatment. The quantifications were achieved by normalization of the peaks to the internal standard geraniol and comparison with the standard curves of authentic 4-methylvaleric acid and 5- methylhexanoic acid (n = 3; means ± SD). EV, empty vector; FW, fresh weight.

To test whether the introduction of *PcIBMS1* or *PcIPMS1* can lead to the production of free 4-methylvaleric acid or an increase in its content, *N. benthamiana* leaf samples were extracted with MTBE, and the MTBE extracts were further analyzed by GC-MS. No free 4-methylvaleric acid was detected in control *N. benthamiana* leaves and in *N. benthamiana* leaves transiently expressing *PcIBMS1* or *PcIPMS1* (Fig. 5b). Since it had been previously observed that compounds with carboxylic groups generated in *N. benthamiana* leaves by the introduction of heterologous genes tend to be modified by endogenous glycosylases (Yang *et al*., 2011; Xu *et al*., 2018), we searched for glycosylated 4-methylvaleric acid by preforming hydrolysis on leaf samples with 0.4N NaOH at 80 ^0^C for 20 min. Surprisingly, 4-Methylvaleric acid and 5-methylhexanoic acid (containing one more carbon than 4- methylvaleric acid) were detected at low levels in the hydrolyzed MTBE extracts of control *N. benthamiana* leaves. However, these two compounds in *N. benthamiana* leaves expressing *PcIBMS1* showed 6.9-fold and 2.7-fold increases in their contents, respectively, compared with that in control *N. benthamiana* leaves (Fig. 5b and c). Transient expression of *PcIPMS1* did not result in any observable changes in the contents of these two branched chain fatty acids in the hydrolyzed MTBE extracts compared with that in the control *N. benthamiana* leaves (Fig. 5b and c). In addition, the relative content of Leu in *N. benthamiana* leaves expressing *PcIBMS1* declined to about 58% compared with that in control *N. benthamiana* leaves (Fig. S3), while no change in Leu concentrations was observed in *N. benthamiana* leaves expressing *PcIPMS1*, which yielded almost the same content of Leu as control *N. benthamiana* leaves (Fig. S3).

### Leu feedback inhibition assay of PcIBMS1 and PcIPMS1

IPMSs from plants and bacteria generally form a homodimeric protein (Koon *et al*., 2004; de Carvalho *et al*., 2005; de Kraker *et al*., 2007; de Kraker & Gershenzon, 2011). Each monomer contains two core domains including an N-terminal catalytic region consisting mainly of a (β/α)8 barrel (TIM barrel) with a divalent metal cofactor necessary for substrate binding (Koon *et al*., 2004), and a C-terminal allosteric regulatory domain responsible for Leu feedback inhibition in microbes and plants (de Carvalho *et al*., 2005; de Kraker & Gershenzon, 2011). The two major domains are separated by two small subdomains that have a flexible hinge in between them (Koon *et al*., 2004). The full-length *PcIBMS1*gene encodes a protein containing 591 amino acids, showing 74%, 66% and 64% amino acid identities with PcIPMS1, AtIPMS1 and OsIPMS1, respectively (Fig. S4), the latter two proteins functionally identified as typical IPMSs in *Arabidopsis thaliana* and *Oryza sativa* (de Kraker *et al*., 2007; Xing & Last, 2017; He *et al*., 2019). Sequence comparison with other typical IPMSs showed that the protein encoded by *PcIBMS1* gene contains the N-terminal catalytic region and likely C-terminal allosteric regulatory domain (Fig. S4). However, many changes in residues occur in the PcIBMS1 protein at positions corresponding to the conserved sites of the other typical IPMS proteins in both N-terminal catalytic region and C-terminal allosteric regulatory domain, notably the loss of five amino acids in the C-terminal domain of the PcIBMS1 protein (Fig. 6 and Fig. S5). The sequence divergence in the C-terminal allosteric regulatory domain of PcIBMS1 protein compared with other typical IPMSs that are sensitive to Leu feedback inhibition led us to hypothesize that *PcIBMS1* encodes an IPMS family protein that is not subject to Leu feedback inhibition.

**Fig. 6.**
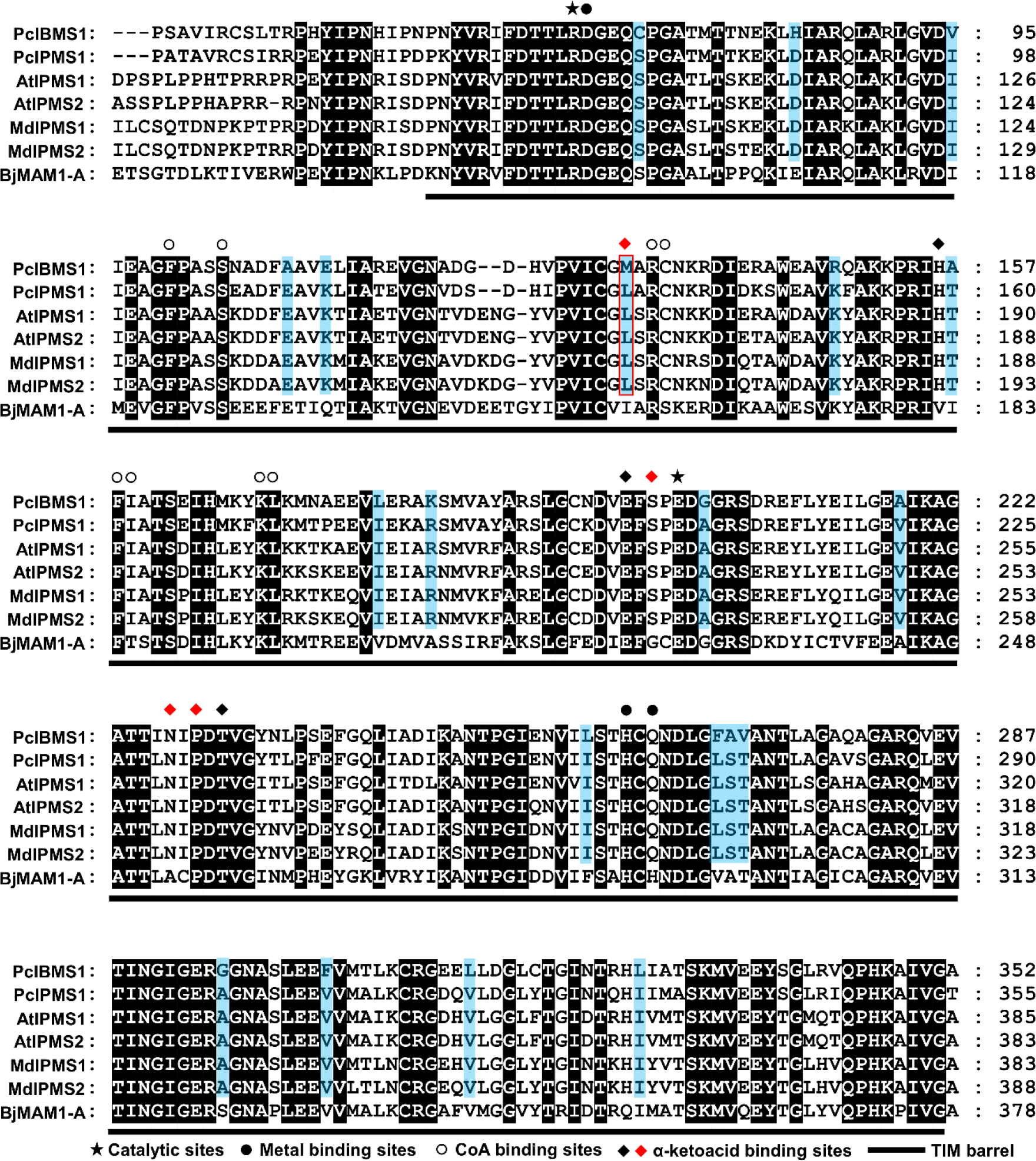
Multiple sequence alignment of N-terminal catalytic regions of *P. cablin*, *M. domestica* and Arabidopsis IPMSs and *B. juncea* MAM1-A. Multiple sequence alignment of the N-terminal catalytic regions of *P. cablin*, *M. domestica* and Arabidopsis IPMSs and BjMAM1-A is shown. Sequence features corresponding to the characterized BjMAM1-A crystal structure are shown (Kumar *et al*., 2019). Residues in catalytic sites, metal binging sites, CoA binding sites and α-ketoacid binding sites are marked by asterisk, sold circle symbol, hollow circle symbol and sold diamond symbol, respectively. The black sold diamond symbol depicts α-ketoacid binding sites implicated in substrate selectivity, while red sold diamond symbol are those shown to impact substrate size discrimination (de Kraker & Gershenzon, 2011; Kumar *et al*., 2019). Protein domain designations are derived from BjMAM1-A crystal structure (Kumar *et al*., 2019). Black background shows perfectly conserved sequences across the IPMS, while the light blue background shows amino acids in the conserved sites in the selected five typical IPMSs diverge from that in the corresponding sites in PcIBMS1. PcIBMS1 and PcIPMS1 were directly obtained from *P. cablin* transcriptome database. The information of other proteins used in this sequence alignment is shown in Table S5.

To test this hypothesis, both recombinant PcIBMS1 and PcIPMS1 were assayed for aldol-type condensation activities between acetyl-CoA and 4-methyl-2-oxovalerate and 2- oxoisovalerate, in the end-point enzyme assay (DTNB) in the presence of Leu at a series of concentrations from 0.05 mM to 10 mM. PcIBMS1 activity was insensitive to Leu concentrations up to the highest concentration measured (10 mM) no matter whether the 2-oxo acid substrate is 4-methyl-2-oxovalerate or 2-oxoisovalerate (Fig. 7a and b). On the other hand, the activity of PcIPMS1 with the substrate 2-oxoisovalerate declined to 57.8% at 50 μM Leu and was reduced to 34.7% at the highest Leu concentration measured (10 mM) (Fig. 7b). The observations of substrate specificity and kinetic properties of PcIPMS1 enzyme towards 2-oxoisovalerate and acetyl-CoA and its sensitivity to Leu feedback inhibition indicate that PcIPMS1 encodes a bona fide IPMS enzyme responsible for Leu biosynthesis.

**Fig. 7.**
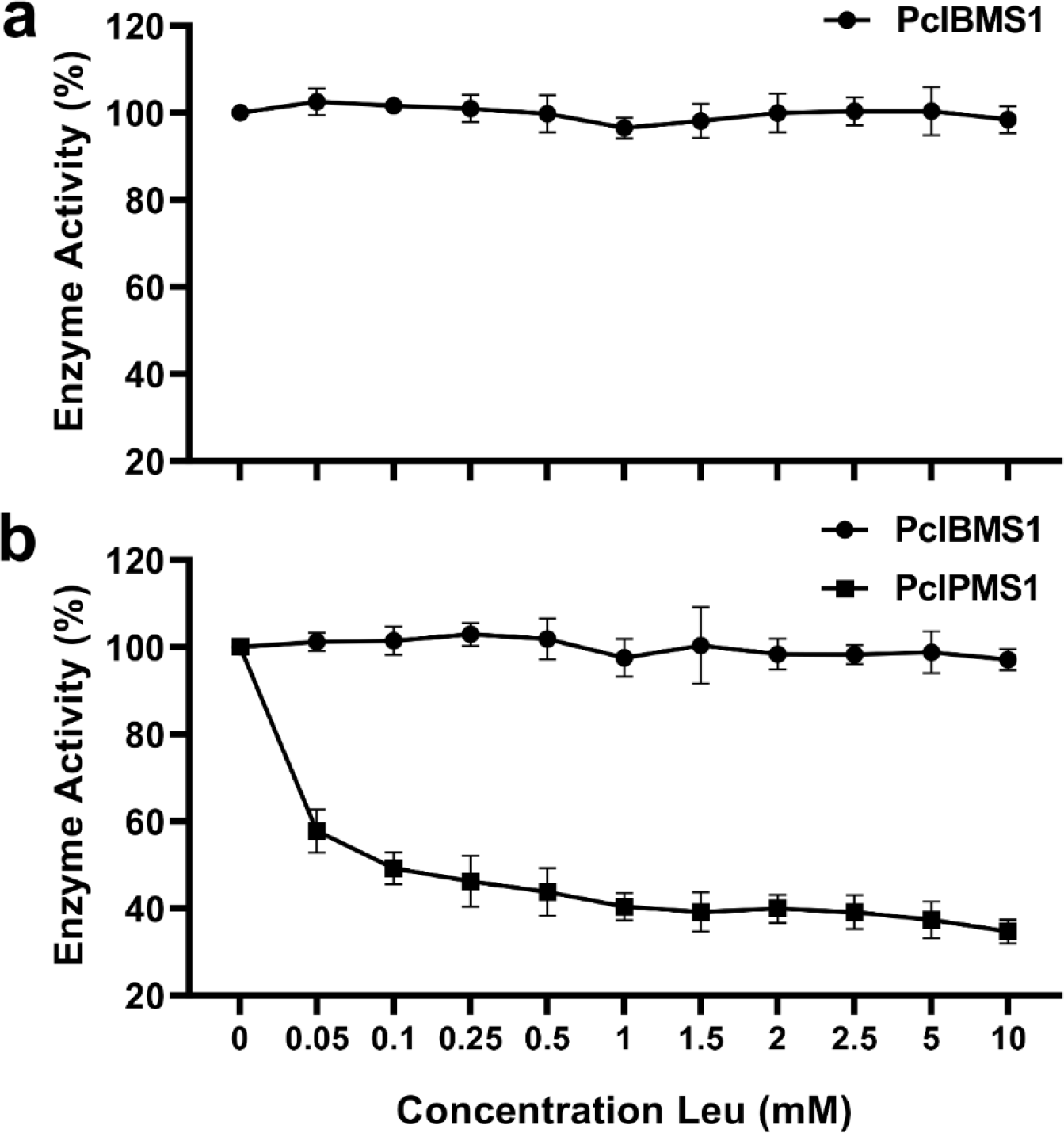
Effect of Leu on the enzymatic activity of PcIBMS1 and PcIPMS1. The effect of Leu on the enzymatic activities of PcIBMS1 towards 4-methyl-2-oxovalerate (**a**) and of both PcIBMS1 and PcIPMS1 towards 2-oxoisovalerate (**b**). Activities are expressed as a percentage of the activity. The 100% activity was set in the absence of Leu. Data represent means ± SD of three independent biological replicates.

### Determination of amino acid sites in the N-terminal domain that may affect the substrate specificity of PcIBMS1 and PcIPMS1

Our data indicates that PcIBMS1 is an IPMS-like protein with an altered preferred substrate (4-methyl-2-oxovalerate rather than 2-oxoisovalerate). Previous studies on the 3D structure of MtIPMS from *Mycobacterium tuberculosis* and BjMAM1-A from *Brassica juncea* identified seven amino acid sites in the N-terminal catalytic domain that may affect the 2-oxo acid substrate chain length specificities of IPMS and methylthioalkylmalate synthase (MAMS) (de Kraker & Gershenzon, 2011; Kumar *et al*., 2019), the latter a neofunctionalized form of IPMS involved in the biosynthesis of Methionine-Derived glucosinolates (de Kraker *et al*., 2007; de Kraker & Gershenzon, 2011). The overall fold of MAMs shares with IPMSs the N-terminal catalytic α/β-barrel domain and the C-terminal α-helical region that forms part of the CoA binding site (Kumar *et al*., 2019). The two major structural differences between MAMs and IPMSs are the loss of the N-terminal extension and the C-terminal Leu binding regulatory domain found in the Leu biosynthesis enzyme. In both MAMs and IPMSs, the catalytic machinery and reaction chemistry are conserved (Kumar *et al*., 2019).

Sequence comparison of the N-terminal domain of PcIBMS1 that catalyzes the extension of the longer chain substrate 4-methyl-2-oxovalerate with those of the five IPMS proteins, including PcIPMS1, AtIPMS1, AtIPMS2, MdIPMS1 and MdIPMS2, all of which show no or negligible activities towards the longer chain substrate 4-methyl-2-oxovalerate compared with that towards the classic IPMS substrate 2-oxoisovalerate (de Kraker *et al*., 2007; Sugimoto *et al*., 2021), highlights the sequence divergence in PcIBMS1 (Fig. 6). Twenty conserved amino acid sites of N-terminal catalytic domains of these five typical IPMS proteins diverged from the corresponding sites in PcIBMS1 protein (Fig. 6). In particular, the conserved Leu located in one of the seven sites in these IPMSs that were identified as accounting for the differences in substrate binding between IPMSs and MAMs (de Kraker & Gershenzon, 2011; Kumar *et al*., 2019), is replaced by Met^132^ in the same position in PcIBMS1 (Fig. 6), suggesting that the substituted Met^132^ may play a vital role in the alteration of substrate specificity of PcIBMS1 from typical IPMSs.

To test the effect of Met^132^ in determining substrate specificity of PcIBMS1, we mutated this residue in PcIBMS1 to the corresponding residue Leu found in the typical IPMS, and we did the reciprocal experiment by changing the corresponding Leu^135^ in PcIPMS1 to Met. We assayed the resulting PcIBMS1 M132L and PcIPMS1 L135M point mutants using 4-methyl-2-oxovalerate and 2-oxoisovalerate and determined their kinetic parameters (Table 3). The PcIBMS1 M132L mutant exhibited about 3-fold decrease in *Km* value for 2-oxoisovalerate with a corresponding approximate 2.4-fold increase in catalytic efficiency (*kcat*/*Km*), while showing only a modest difference in the *Km* value for 4-methyl- 2-oxovalerate and about 2-fold decrease in catalytic efficiency versus the wild type. The PcIPMS1 L135M point mutant exhibited about 2.2-fold increase in *Km* value for 2- oxoisovalerate with a corresponding approximate 5-fold decrease in catalytic efficiency (*kcat*/*Km*) versus the wild type, while still showing no detectable activities with 4-methyl-2- oxovalerate. These results suggest that this amino acid site indeed contributes to the ability of the enzyme to accept the longer chain 2-oxo acid 4-methyl-2-oxovalerate but that other amino acids play a crucial role as well.

### Phylogenetic relationships of PcIBMS1 homologs

To gain further insight into the evolution of PcIBMS1, we expanded the phylogenetic analysis of PcIBMS1 and PcIPMS1 to include their closest homologs from plant species and microorganisms, including bona fide IPMSs, MAMs from *Arabidopsis thaliana* and *Brassica juncea*, and citramalate synthase (CMS) from *Malus domestica* (Fig. 8). PcIBMS1, MdCMS1 and MAMs, as well as SlIPMS3, a tomato true IPMS that is nonetheless lacking the regulatory C-terminal regulatory domain and is involved in synthesizing the C5 acyl precursor for acylsugar biosynthesis rather than leucine, split off from typical plant IPMSs prior to the split between monocot and dicot IPMSs (Fig. 8a). However, these leucine-insensitive enzymes represent neofunctionalized forms of typical plant IPMS (Ning *et al*., 2015; Kumar *et al*., 2019; Sugimoto *et al*., 2021), that either lack the regulatory C-terminus domain altogther or possess the C-terminal domain but show high rate of change in it. We therefore redid the phylogenetic analysis with the protein sequences from which the C-terminal regulatory domain was removed (Fig. 8b). The phylogenetic tree obtained in this analysis showed that PcIBMS1 is most closely related to PcIPMS1.

**Fig. 8.**
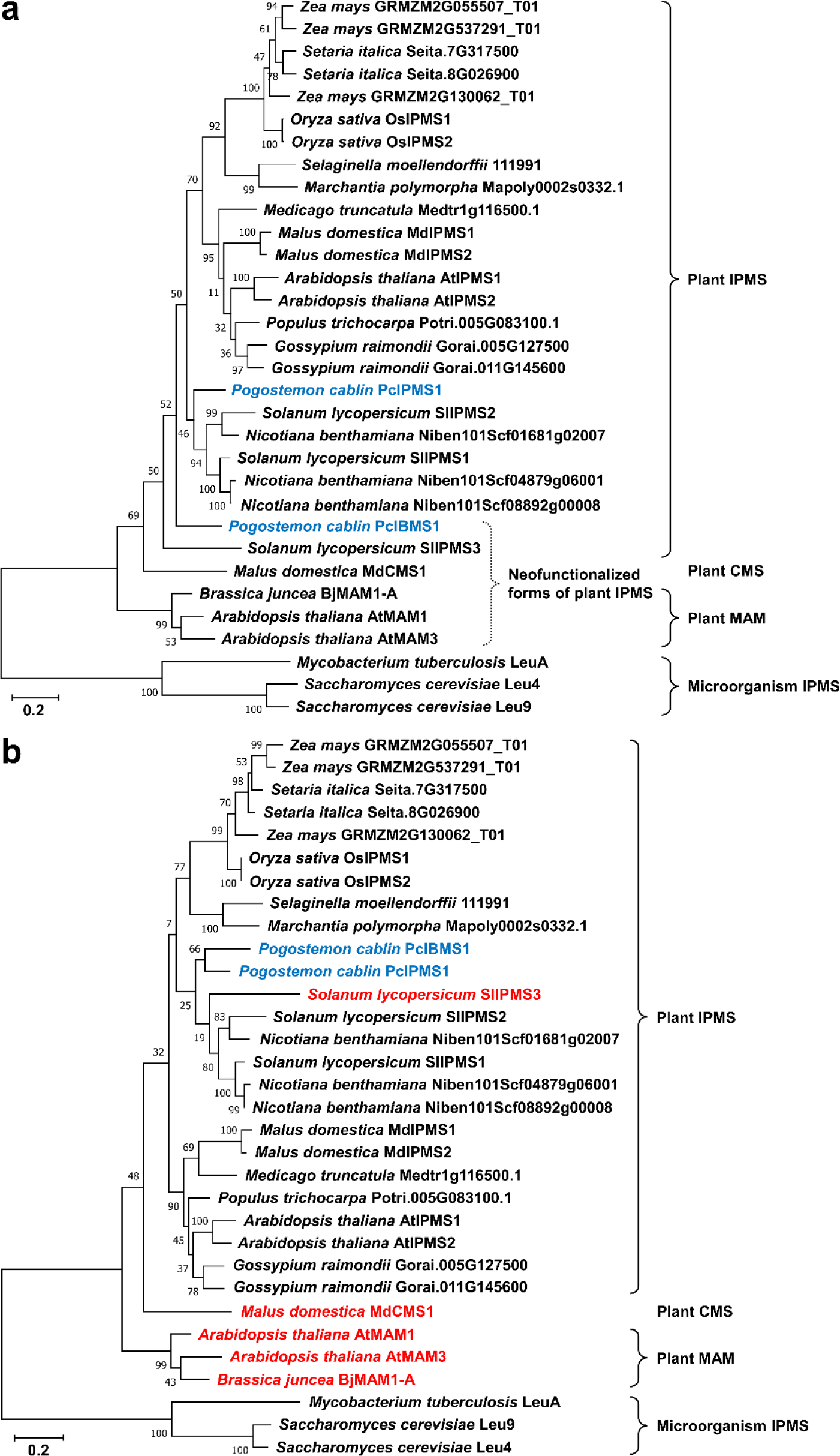
Phylogenetic analyses of PcIBMS1 homologs, methylthioalkymalate synthases (MAMSs) and citramalate synthase (CMS) from selected species. **a.** Phylogenetic analyses of whole PcIBMS1 homologs, MAMSs and CMS from selected species. The whole protein sequences of PcIBMS1 homologs, MAMSs and CMS were used in this phylogenetic analysis. **b.** Phylogenetic analyses of the N-terminal domains of PcIBMS1 homologs, MAMSs and CMS from selected species. PcIBMS1 and all the IPMSs except SlIPMS3, from which the predicted C-terminal allosteric regulatory domains were removed, were used in this phylogenetic analysis. The proteins naturally lacking the C- terminal allosteric regulatory domain that is presented in typical IPMSs were marked in red color. For the construction of phylogenetic trees in (**a**) and (**b**), all proteins except PcIBMS1 and PcIPMS1 were obtained from the PHYTOZONE v12.1 (https://phytozome.jgi.doe.gov/pz/portal.html), NCBI website (https://www.ncbi.nlm.nih.gov/), TAIR website (https://www.arabidopsis.org/) and UniProt website (https://www.uniprot.org/) (Table S5). PcIBMS1 and PcIPMS1, the protein sequences of which were directly obtained from *P. cablin* transcriptome database, were marked in blue color. Sequences were aligned using MUSCLE (Edgar, 2004), and the trees were conducted using the Maximum likelihood algorithm and JTT matrix-based model in MEGA X (Jones *et al*., 1992; Kumar *et al*., 2018). Branch point bootstrap values were calculated with 1000 replicates. The percentage of trees in which the associated taxa clustered together is shown next to the branches. The tree is drawn to scale, with branch lengths measured in the number of substitutions per site. The species were listed along with the protein names.

## DISCUSSION

### 4-Methylvaleric acid is a precursor of pogostone

Studies have shown that pogostone, one of the major chemical components of patchouli oil from *Pogostemon cablin*, possesses many pharmaceutical activities and thus could be considered as a potential therapeutic agent for treatment of many diseases. In our previous study, we constructed a *P. cablin* RNAseq database, from which we isolated and characterized PcAAE2, an enzyme that catalyzes the formation of 4-methylvaleryl-CoA from 4-methylvaleric acid (Chen, J *et al*., 2021). The expression of *PcAAE2* correlates spatially and temporally with pogostone biosynthesis, and based on this data as well as the structure of pogostone, we further hypothesized that 4-methylvaleric acid is a precursor of pogostone (via 4-methylvaleryl-CoA). Here, by feeding *P. cablin* young leaves with deuterium-labeled [^2^H11] 4-methylvaleric acid (4MVA-d11) and following the incorporation of this label into pogostone, we were able to show that 4-methylvalerate is indeed a precursor in pogostone biosynthesis (Fig. 2).

### PcIBMS1 catalyzes the formation of 4-methylvaleric acid

To further elucidate the route leading to biosynthesis of pogostone, we attempted to figure out the key enzymes involved in the biosynthesis of 4-methylvaleric acid. This branched C6 fatty acid has a similar structure to that of 3-methylbutanoic acid, except that it has one more carbon than the latter. The synthesis of 3-methylbutanoic acid, a Leu precursor, is well documented and involves the α-ketoacid elongation (αKAE) (Binder, 2010). Therefore, we predicted that 4-methylvaleric acid is derived from leucine metabolism by one additional elongation cycle through αKAE pathway.

Gene candidates for 4-methylvaleric acid biosynthesis were identified by utilizing Arabidopsis representative genes involved in leucine metabolism to query the assembled *P. cablin* transcriptome data through homology search by TBLASTN software. A total of 13 unique genes were obtained. One of these unique genes, designated as *PcIBMS1*, showed the highest correlation with pogostone biosynthesis markers and was homologous to IPMS, the key enzyme in the one-carbon elongation of 2-oxoisovalerate to 4-methyl-2- oxovalerate (via 2-isopropylmalate) in Leu biosynthesis (Fig. 1, Table 1 and Fig. S1).

Based on the identification of *PcIBMS1* as the most likely candidate encoding the key enzyme for 4-methylvaleric acid biosynthesis, we proceeded to produce and purify the recombinant PcIBMS1 protein and test its catalytic activities in *in vitro* biochemical assays (Table 2, Table 3 and Fig. 4). In these assays, PcIBMS1 effectively catalyzed the formation of 2-isobutylmalate from 4-methyl-2-oxovalerate and acetyl-CoA. In addition, PcIBMS1 was also able to catalyze the formation of 2-isopropylmalate from 2-oxoisovalerate and acetyl-CoA.

Analysis of compounds produced in *N. benthamiana* leaves when *PcIBMS1* was transiently expressed lent additional evidence for its function in 4-methylvaleric acid biosynthesis. We were able to directly detect free 2-isopropylmalate and 2-isobutylmalate, which were not produced in the control, as well as a large increase in the content of glycosylated 4-methylvaleric acid and glycosylated 5-methylhexanoic acid (Fig. 5). These results indicate that when *PcIBMS1* is overexpressed in *N. benthamiana*, the resulting protein can also act with 2-oxoisovalerate, despite its high *Km* value for this substrate, and also with 5-methyl-2-oxohexanoate. It further indicates that endogenous *N. benthamiana* proteins can complete the elongation cycle with 2-isobutylmalate and longer intermediates.

*N. benthamiana* is known to synthesize such longer branched chain fatty acids as part of its acylsugar biosynthesis in glandular trichomes (Kroumova & Wagner, 2003; Slocombe *et al*., 2008), which is a possible explanation why we detected low levels of glycosylated 4-methylvaleric acid and glycosylated 5-methylhexanoic acid in control *N. benthamiana* leaves.

We also isolated and characterized a second IPMS-like gene from *P. cablin* whose expression pattern did not correlate with pogostone biosynthesis. The protein encoded by this gene, designated as PcIPMS1, was characterized in a similar way and was shown to be an effective isopropylmalate synthase using 2-oxoisovalerate as a substrate, with kinetic properties similar to other plant IPMSs (Table 3; de Kraker *et al*., 2007; Ning *et al*., 2015). It did not have activity with 4-methyl-2-oxovalerate (Table 3). The *Km* value of PcIBMS1 for 2-oxoisovalerate was almost 5-fold higher than that of PcIPMS1, but its *Km* value for 4-methyl-2-oxovalerate was 3-fold lower than the *Km* value of PcIPMS1 with 2- oxoisovalerate. However, PcIBMS1 has a somewhat high *Km* value for acetyl-CoA when acting on its preferred substrate, compared with IPMSs (Table 3; de Kraker *et al*., 2007; Ning *et al*., 2015).

### Met^132^ amino acid site in PcIBMS1 affects its substrate preference

MAM structure of a MAMS from *Brassica juncea*, designated as BjMAM1-A, was determined by X-ray crystallography (Kumar *et al*., 2019). Seven amino acid sites that may affect substrate preference of IPMS and MAMS were identified. Our sequence comparisons showed that one of the seven corresponding sites is substituted by Met^132^ in PcIBMS1 (Fig. 6). This site corresponds to Leu^143^ in Mt-IPMS that was shown to have hydrophobic interaction with one of the two methyl groups of 2-oxoisovalerate by RCSB PDB Ligand Explorer 3.5 (Koon *et al*., 2004; de Kraker & Gershenzon, 2011). Therefore, we hypothesized that Met^132^ may play a vital role in substrate preference of PcIBMS1. PcIBMS1 M132L mutant obtained by site-directed mutagenesis showed much higher affinity and catalytic efficiency towards 2-oxoisovalerate and lower catalytic efficiency towards 4-methyl-2-oxovalerate than wild-type PcIBMS1 (Table 3). The reciprocal PcIPMS1 mutant, PcIPMS1 L135M, showed a strong decrease in affinity and catalytic efficiency towards 2-oxoisovalerate compared with the wild-type enzyme, but it still did not have the capability of utilizing the larger substrate 4-methyl-2-oxovalerate. Therefore, while Met^132^ plays a critical role in PcIBMS1 substrate specificity, simply substituting Leu in this position in IPMS, is clearly not sufficient to convert it into IBMS.

### PcIBMS1 has altered feedback regulation of enzyme activity

PcIPMS1 is a typical IPMS enzyme which exhibits Leu feedback inhibition both in vitro (Fig. 7) and in planta (overexpression of *PcIPMS1* in *N. benthamiana* did not lead to an increase in Leu concentrations, Fig. S3). In contrast, our sequence comparisons showed that PcIBMS1 has a wide sequence divergence from typical plant IPMS proteins in both the catalytic N-terminal and the C-terminal regulatory domain (Fig. 6 and Fig. S5). In the latter domain, a deletion of five amino acids in PcIBMS1 was particularly notable (Fig. S5). Feedback inhibition assay indicated that *PcIBMS1* encodes an enzyme that is not subject to Leu inhibition (Fig. 7). The loss of Leu regulation by the sequence divergence and the loss of five amino acids in PcIBMS1 were thus obtained by a different mechanism of the complete loss of the C-terminal domain, which is how Brassica MAMS and Solanaceae SlIPMS3 acquired their insensitivity to Leu inhibition. Further structure analysis and gain-of-function mutations in the C-terminal regulatory domain of PcIBMS1 might provide a better understanding of the mechanism leading to the insensitivity to feedback inhibition of this enzyme.

### IPMSs have been recruited and neofunctionalized to specialized metabolism pathways multiple times throughout plant evolution

Several instances of IPMS neofunctionalization and recruitment to specialized metabolism within eudicots have been documented, including Brassicaceae MAMSs involved in biosynthesis of Met-related glucosinolate compounds, apple CMS involved in flavor esters, and wild tomato *IPMS3* involved in acylsugar biosynthesis (Ning *et al*., 2015; Kumar et al., 2019; Sugimoto et al., 2021). In these three cases, a loss of the C-terminal allosteric regulatory domain has occurred, which freed these enzymes from Leu inhibition. When the phylogeny of these enzymes is analyzed by standard methodology, they appear to have split from the bona fide IPMS clade before the split of monocot and dicot IPMSs (Fig. 8a). However, this analysis is probably misleading, since the genes encoding these enzymes have undergone a major change involving the loss of a domain and likely accelerated substitutions elsewhere. When only the catalytic domain is used in the phylogenetic analysis, MAMs and CMS still split early from the IPMS clade, but PcIBMS1 now clusters with PcIPMS1, and *wild tomato* IPMS3 groups with other Solanaceae IPMSs (Fig. 8b). This analysis indicates that at least in the case of PcIBMS1, its recruitment and neofunctionalization into the pogostone pathway occurred relatively recently.

Besides the changes in the catalytic domain that are obviously necessary to evolve a new substrate specificity, the uniform loss of Leu inhibition is of interest. Since the end product of the pathways that these enzymes participate in is not Leu, a Leu sensitivity will serve no clear function. But perhaps more importantly, specialized metabolites such as glucosinolates, flavor esters, acylsugars, and pogostone are typically made in large quantities in specific tissues or organs, and being inhibited by related metabolites would be detrimental to achieving such high levels of production.

Based on our results with PcIBMS and the previous research on the origin of MAMs and CMS, it is likely that many more IPMS-derived enzymes responsible for synthesizing substrates for the biosynthesis of specialized metabolism are still to be discovered. In particular, one anticipates such genes in the Solanaceae, where many species synthesize acyl sugars using acyls up to 12 carbons long, and where the αKAE pathway was shown to be involved in *Nicotiana* and *Petunia*. In fact, our detection in *N. benthamiana* leaves (not expressing PcIBMS) of basal levels of 4-methylvaleric acid and 5-methylhexanoic acid (we did not look at longer acids) suggests that *N. benthamiana* has endogenous enzymes related to IPMS/IBMS. Furthermore, there are three IPMS/IBMS homologs identified in the *N. benthamiana* genome (Fig. 8), and it thus remains to be determined if one or more of these could be involved in the synthesis of 4-methylvaleric acid and 5- methylhexanoic acid and other related but longer acids.

## Data availability

Data supporting the finding of this study are available within the article and its supplemental files. The sequence data used in this study can be obtained from NCBI with the following GenBank accession numbers: PcIBMS1, MW413958; PcIPMS1, MW413959.

## Acknowledgements

We would like to thank Dr. Deng at Analytical and Testing Center of Chongqing University for their assistance with LC-QTOF-MS analysis and the laser scanning confocal microscopy analysis.

## Author contributions

H.X. designed the experiments; C.W., Y.W., J.C., L.L. and Z.L. conducted the experiments, analyzed data or provided material; E.P. and H.X wrote the article; all authors edited the article.

## Supporting Information

Fig. S1 RT-qPCR analysis of transcript levels of gene candidates involved in 4- methylvaleric acid biosynthesis in different tissues of *Pogostemon cablin* at different stages of development.

Fig. S2 Dependency of PcIBMS1 and PcIPMS1 enzymatic activity on incubation conditions and components.

Fig. S3 LC-QTOF-MS analyses of relative abundances of leucine in *N. benthamiana* leaves expressing *PcIBMS1* and *PcIPMS1*.

Fig. S4 Sequence alignment of PcIBMS1, PcIPMS1, AtIPMS1, OsIPMS1, SlIPMS3 and MtIPMS.

Fig. S5 Sequence alignment of C-terminal domain of PcIBMS1, PcIPMS1, MtIPMS and other plant typical IPMSs that are subject to Leu feedback inhibition.

Table S1 All primers used in this present study.

Table S2 Bioinformatic analysis of gene candidates involved in 4-methylvaleric acid biosynthesis screened from *P*. *cablin* RNAseq database.

Table S3 Average normalized counts of gene candidates involved in 4-methylvaleric acid biosynthesis and *PcAAE2*in *P. cablin* RNA-seq database.

Table S4 Ranking of the top 10 unique genes in the *P. cablin* RNAseq database by coexpression analysis with the *PcAAE2* gene.

Table S5 GenBank accession or locus numbers of functional IPMSs, MAMs and MdCMS1 from NCBI (plants), TAIR and UniProt (bacteria and yeast) sites used for phylogenetic reconstruction and sequence comparison.

