## Supporting information for "Synthesis of 4-methylvaleric acid, a precursor of pogostone, involves a 2-isobutylmalate synthase related to 2-isopropylmalate synthase of leucine biosynthesis"

**Table S1 All primers used in this present study**

| Name of primers | DNA sequences (5'- 3') |
|---|---|
| **Vector construction for heterologous expression in *E. coli*** | |
| Construction of pET-28a-PcIPMS1 and pET-28a-PcIPMS2 | |
| PcIBMS1-TRTN-F | TGGTGCCGCGCGGCAGCCATATGTCCCTCACTCGCCCTCACTACATC |
| PcIBMS1-PETN-R | GTCGACGGAGCTCGAATTCGTCATTTCAAATCAAGCAACGTCTCATTG |
| PcIPMS1-TRTN-F | TGGTGCCGCGCGGCAGCCATATGGTCCGCTGTTCCATCCG |
| PcIPMS1-PETN-R | GTCGACGGAGCTCGAATTCGTTACTGGAATGCATTTACCACCTTACTTTCTTG |
| Construction of pET-28a-PcIPMS1 M132L | |
| PcIBMS1-M132L-F1 | TGGTGCCGCGCGGCAGCCATATGTCCCTCACTCGCCCTCAC |
| PcIBMS1-M132L-R1 | GAGGCCGCAGATCACCGGC |
| PcIBMS1-M132L-F2 | TGCCGGTGATCTGCGGCCTC**GCG**AGGTGCAACAAGAGGG |
| PcIBMS1-M132L-R2 | GTCGACGGAGCTCGAATTCGTCATTTCAAATCAAGCAACGTCTC |
| Construction of pET-28a-PcIPMS2 L135M | |
| PcIPMS1-L135M-F1 | TGGTGCCGCGCGGCAGCCATATGGTCCGCTGTTCCATCCG |
| PcIPMS1-L135M-R1 | TCCACAGATCACCGGGATATG |
| PcIPMS1-L135M-F2 | CATATCCCGGTGATCTGTGGA**ATG**GCGAGGTGTAATAAGAGGGATATTG |
| PcIPMS1-L135M-R2 | GTCGACGGAGCTCGAATTCGTTACTGGAATGCATTTACCACCTTAC |
| **Vector construction for transient heterologous expression in *Nicotiana benthamiana*** | |
| PcIBMS1-AQ-F | ATATTCTGCCCAAATTCGCGAATGGCGTCTTCCTCCTCTTCC |
| PcIBMS1-AQ-R | ATGAAACCAGAGTTAAAGGCCTCATTTCAAATCAAGCAACGTCTC |
| PcIPMS1-AQ-F | ATATTCTGCCCAAATTCGCGAATGGCGACATCCATTTGCAAG |
| PcIPMS1-AQ-R | ATGAAACCAGAGTTAAAGGCCTTACTGGAATGCATTTACCACCTTAC |
| **For RT-qPCR analysis** | |
| PcIBMS1-RT-F | CGAAGAAGCCGAGGATTCAC |
| PcIBMS1-RT-R | CTCTTAGCCCTCTCCAACACC |
| PcIPMS1-RT-F | GTCTCCTGAAGATGTTGGACTTC |
| PcIPMS1-RT-R | AATGGCCTTGAAGCGTGTG |
| PcIPMILSSU1-RT-F | GGATTGCCCTCCACATACCA |
| PcIPMILSSU1-RT-R | GAGCCGCACCCAAAGTTATC |
| PcIPMILLSU1-RT-F | CGCAAACCTTCGACTACTGG |

| | |
|---|---|
| PcIPMILLSU1-RT-R | TGTGGTTTCTCTGACGCTCT |
| PcIPMDHL1-RT-F | TGTTGGAGGCTTCCATGCTA |
| PcIPMDHL1-RT-R | CAAACTGCTTTGGGTTTCGC |
| BCKDHL-E1α-1- RT-F | CCTGGTGGAAGGGTCTCATT |
| BCKDHL-E1α-1- RT-R | TGAAGGGTCACCATGCTTCT |
| BCKDHL-E1α-2- RT-F | CACTTGCTGTGTTCAGTGCT |
| BCKDHL-E1α-2- RT-R | AACTGCTCTCCACCATTCCA |
| BCKDHL-E1β- RT-F | TGCTGAGATTTCTGCCTCCA |
| BCKDHL-E1β- RT-R | ATGGCTCGAACACAAGAGGA |
| BCKDHL-E2- RT-F | CAGTCGCTCTCAATCTCGGA |
| BCKDHL-E2- RT-R | CTTCCCACCAATTGCACCAA |
| BCKDHL-E3- RT-F | TGGATGGTGAAGTCCGGAAA |
| BCKDHL-E3- RT-R | CATCTGCCTCTAGGGTGGTC |
| PcBCATL1- RT-F | ACATTGCACGAGATGAAGGC |
| PcBCATL1- RT-R | GCTTCCTACAGGAGCAACAC |
| PcBCATL2- RT-F | CATCGCCTCGTAGCAATCAC |
| PcBCATL2- RT-R | TGAACACCTCCTCGTCTTCC |
| PcBCATL3- RT-F | TGAGCCCATCAGCTGGTATT |
| PcBCATL3- RT-R | CACCCACCCTCAGTCTCATT |
| PcGAPDH-RT-F | GATTGGAGAGGTGGACGAGC |
| PcGAPDH-RT-R | CAACAGTTGGGACACGGAAG |
| PcActin7-RT-F | TACAGAGGCACCACTCAACC |
| PcActin7-RT-R | CGACCACTAGCGTACAGTGAG |
| PcTubulin3-RT-F | GCCTGGACAGAGTGAGGAAG |
| PcTubulin3-RT-R | TCCATAATCAACCGACAACC |
| **Vector construction for subcellular localization analysis** | |
| PTF-PcIBMS1-F | TTACTATTTACAATTACAGTCGACATGGCGTCTTCCTCCTCTTCCCTC |
| PTF-PcIBMS1-R | CTCGCCCTTGCTCACCATGGATCCTTTCAAATCAAGCAACGTCTCATTGTG |
| PTF-PcIPMS1-F | TTACTATTTACAATTACAGTCGACATGGCGACATCCATTTGCAAGCAC |
| PTF-PcIPMS1-R | CTCGCCCTTGCTCACCATGGATCCCTGGAATGCATTTACCACCTTACTTTCTTGG |

The overlaps (15–30 bp) in the primer sequences that were used for DNA fragment assembly were underlined. Substituted sequences for construction of PcIBMS1 M132L and PcIPMS1 L135M point mutants are marked in bold and red color.

**Table S2 Bioinformatic analysis of gene candidates involved in 4-methylvaleric acid biosynthesis screened from *P. cablin* RNAseq database.**

| Genes | Closest Arabidopsis homolog, accession no. and amino acid identity | Functional annotation of the closest Arabidopsis homolog | Peptide length (amino acids) | Signal peptide prediction (using Target P-2.0 and PTS1 Predictor) |
|---|---|---|---|---|
| PcIBMS1 | AT1G74040 (71%) | 2-Isopropylmalate synthase 1 | 591 | mTP (0.0013), **cTP (0.9729)**, luTP (0.0254), PTS1 (-48.477) |
| PcIPMS1 | AT1G74040 (78%) | 2-Isopropylmalate synthase 1 | 604 | mTP (0.0013), **cTP (0.9926)**, luTP (0.0024), PTS1 (-41.119) |
| PcIPMIL SSU | AT2G43090 （68%） | Isopropylmalate isomerase small subunit 1 | 255 | mTP (0), **cTP (0.9114)**, luTP (0.0869), PTS1 (-50.713) |
| PcIPMIL LSU | AT4G13430 （81%） | Isopropyl malate isomerase large subunit 1 | 510 | mTP (0.008), **cTP (0.9572)**, luTP (0.0005), PTS1 (-51.337) |
| PcIPMDHL | AT1G80560 （80%） | Isopropylmalate dehydrogenase 2 | 404 | mTP (0.0107), **cTP (0.9699)**, luTP (0.0142), PTS1 (-16.432) |
| PcBCATL1 | AT1G10070 （70%） | Branched-chain amino acid transferase 2 | 392 | **mTP (0.9813)**, cTP (0.0015), luTP (0), PTS1 (-54.105) |
| PcBCATL2 | AT1G10070 （62%） | Branched-chain amino acid transferase 2 | 385 | **mTP (0.7027)**, cTP (0.0012), luTP (0.0001), PTS1 (-53.954) |
| PcBCATL3 | AT5G65780 （78%） | Branched-chain amino acid aminotransferase 5 | 418 | mTP (0.0001), **cTP (0.7168)**, luTP (0.0341), PTS1 (-41.206) |
| PcBCKDHL E1α1 | AT1G21400 （62%） | Thiamin diphosphate-binding fold (THDP-binding) superfamily protein | 482 | **mTP (0.7533)**, cTP (0.0068), luTP (0.0002), PTS1 (-51.494) |
| PcBCKDHL E1α2 | AT5G09300 （67%） | Thiamin diphosphate-binding fold (THDP-binding) superfamily protein | 468 | mTP (0.272), **cTP (0.5969)**, luTP (0.0059), PTS1 (-41.250) |
| PcBCKDHL E1β | AT1G55510 （88%） | Branched-chain alpha-keto acid decarboxylase E1 beta subunit | 347 | **mTP (0.9906)**, cTP (0.0001), luTP (0), PTS1 (-12.473) |
| PcBCKDHL E2 | AT3G06850 （61%） | Dihydrolipoamide branched chain acyltransferase | 523 | **mTP (0.5057)**, cTP (0.0455), luTP (0.0003), PTS1 (-48.029) |
| PcBCKDHL E3 | AT1G48030 （84%） | Mitochondrial lipoamide dehydrogenase 1 | 506 | **mTP (0.9999)**, cTP (0.0001), luTP (0), PTS1 (-25.218) |

The most likely organelle that gene is targeted to and the corresponding prediction value were highlighted in bold.

**Table S3 Average normalized counts of gene candidates involved in 4-methylvaleric acid biosynthesis and *PcAAE2* in *P. cablin* RNA-seq database.**

| **Unigene ID** | **Seeding** | **5W-Root** | **5W-Stem** | **5W-TLeaf** | **8W-TLeaf** | **8W-SLeaf** |
|---|---|---|---|---|---|---|
| TRINITY_DN7984_c0_g1 (PcIBMS1) | 1898.7 | 644.5 | 1672.67 | 1334.5 | 2.3333 | 4.3333 |
| TRINITY_DN5555_c0_g1 (PcIPMS1) | 1722.33 | 1939 | 2364.66 | 1770.3 | 1563.66 | 1980.33 |
| TRINITY_DN5796_c0_g1 (PcIPMIL SSU) | 3597.67 | 2497 | 3380.33 | 3993.5 | 2791 | 2745 |
| TRINITY_DN22473_c0_g3 (PcIPMIL LSU) | 4277 | 2992 | 4018.67 | 5021.5 | 2855 | 4303 |
| TRINITY_DN28440_c0_g2 (PcIPMDHL) | 2537 | 1982.5 | 2675.33 | 2671.5 | 1638.67 | 1346.33 |
| TRINITY_DN3284_c1_g1 (PcPcBCATL1) | 27.3333 | 263.5 | 32.3333 | 38 | 121.997 | 2702.92 |
| TRINITY_DN20752_c0_g2 (PcBCATL2) | 274 | 65 | 208 | 405 | 265 | 132 |
| TRINITY_DN8015_c0_g3 (PcBCATL3) | 1586 | 1753.49 | 1701 | 2119.5 | 1522 | 1220.67 |
| TRINITY_DN6515_c0_g1 (PcBCKDHL E1α1) | 1162 | 685 | 1961.67 | 1182.5 | 1766.67 | 3290.66 |
| TRINITY_DN8285_c0_g2 (PcBCKDHL E1α2) | 705 | 1224.5 | 552.333 | 631.5 | 545 | 605.333 |
| TRINITY_DN12512_c0_g3 (PcBCKDHL E1β) | 1259 | 1335 | 1477.33 | 1234.5 | 1201.33 | 1563 |
| TRINITY_DN9874_c0_g2 (PcBCKDHL E2) | 1070.67 | 2207 | 937.667 | 738 | 654 | 1225.67 |
| TRINITY_DN22140_c0_g2 (PcBCKDHL E3) | 5061.33 | 6519.5 | 3216.67 | 6105.5 | 2386.67 | 3396.67 |
| TRINITY_DN3370_c0_g1 (PcAAE2) | 801.67 | 350 | 974 | 658 | 152.667 | 96 |

**Table S4 Ranking of the top 10 unique genes in the *P. cablin* RNAseq database by coexpression analysis with the *PcAAE2* gene.**

| **Unigene ID** | **Functional annotation by SWISS-PROT database** | **Pearson Correlation Coefficient** | ***P* (Two Tailed)** |
|---|---|---|---|
| TRINITY_DN6205_c0_g3 | ABC transporter B family member 25 | 0.979 | 0.001 |
| TRINITY_DN1882_c0_g1 | Transcription factor | 0.969 | 0.001 |
| **TRINITY_DN7984_c0_g1 (PcIBMS1)** | **2-isopropylmalate synthase A** | **0.969** | **0.001** |
| TRINITY_DN27812_c0_g1 | Unknown protein | 0.965 | 0.002 |
| TRINITY_DN8346_c0_g1 | BTB/POZ domain-containing protein | 0.955 | 0.003 |
| TRINITY_DN26973_c0_g1 | Receptor like protein 29 | 0.955 | 0.003 |
| TRINITY_DN4920_c0_g1 | Protein HOTHEAD | 0.954 | 0.003 |
| TRINITY_DN19765_c0_g1 | Ferritin-3 | 0.954 | 0.003 |
| TRINITY_DN9171_c0_g3 | Histidinol dehydrogenase | 0.953 | 0.003 |
| TRINITY_DN7580_c0_g1 | Mitogen-activated protein kinase | 0.952 | 0.003 |

**Table S5 GenBank accession or locus numbers of functional IPMSs, MAMs and MdCMS1 from NCBI (plants), TAIR and UniProt (bacteria and yeast) sites used for phylogenetic reconstruction and sequence comparison.**

| Sequence name | Source | GenBank accession or locus no. | References |
|---|---|---|---|
| AtIPMS1 | *Arabidopsis thaliana* | AT1G18500 | (de Kraker *et al.*, 2007) |
| AtIPMS2 | | AT1G74040 | |
| AtMAM1 | | AT5G23010 | (Textor *et al.*, 2004) |
| AtMAM3 | | AT5G23020 | (Textor *et al.*, 2007) |
| BjMAM1-A | *Brassica juncea* | CAQ56040 | (Kumar *et al.*, 2019) |
| MdCMS1 | *Malus domestica* | MD05G1155100 | (Sugimoto *et al.*, 2021) |
| MdIPMS1 | | MD15G1332600 | |
| MdIPMS2 | | MD01G1019700 | |
| OsIPMS1 | *Oryza sativa* | Os11g04670.1 | (He *et al.*, 2019) |
| OsIPMS2 | | Os12g04440.1 | |
| SlIPMS1 | *Solanum lycopersicum* | Solyc06g053400.2.1 | (Ning *et al.*, 2015) |
| SlIPMS2 | | Solyc08g014130.2.1 | |
| SlIPMS3 | | Solyc08g014230.2.1 | |
| MtIPMS (LeuA) | *Mycobacterium tuberculosis* H37Rv | P9WQB3 | (Koon *et al.*, 2004) |
| Leu4 | *Saccharomyces cerevisiae* | YNL104C | (Chang *et al.*, 1984) |
| Leu9 | | YOR108W | (Casalone *et al.*, 2000) |

Protein sequences from PHYTOZONE v12.1 website were not included in this table and can be obtained directly from relevant websites through the gene names displayed in the phylogenetic tree.

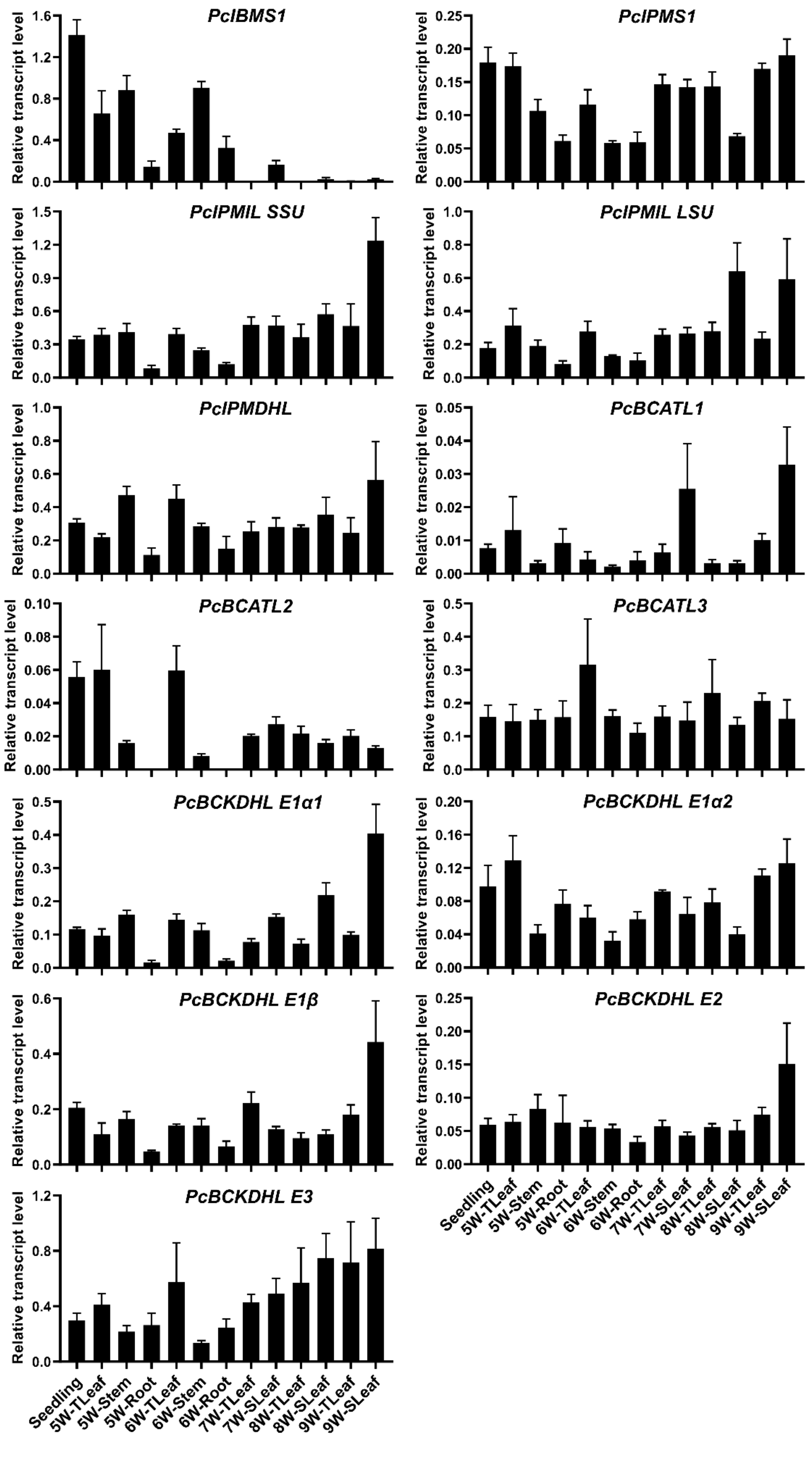
PcIBMS1
PcIPMS1
PcIPMIL SSU
PcIPMIL LSU
PcIPMDHL
PcBCATL1
PcBCATL2
PcBCATL3
PcBCKDHL E1α1
PcBCKDHL E1α2
PcBCKDHL E1β
PcBCKDHL E2
PcBCKDHL E3
Relative transcript level
Seedling
5W-TLeaf
5W-Stem
5W-Root
6W-TLeaf
6W-Stem
6W-Root
7W-TLeaf
7W-SLeaf
8W-TLeaf
8W-SLeaf
9W-TLeaf
9W-SLeaf

39

**Fig. S1 RT-qPCR analysis of transcript levels of gene candidates involved in 4-methylvaleric acid biosynthesis in different tissues of *Pogostemon cablin* at different stages of development.** The RT-qPCR data were presented as relative expression of these genes relative to *P. cablin GAPDH* (glyceraldehyde-3-phosphate dehydrogenase), *Actin 7* and *Tubulin 3* in each tissue sample. Data are means ± SD of four independent biological replicates.

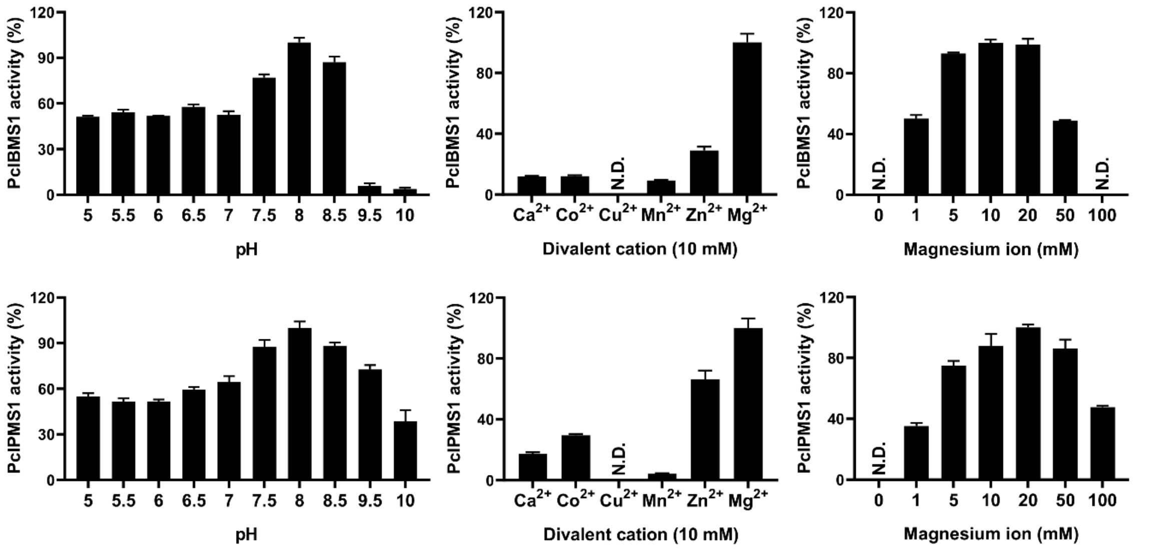


**Fig. S2 Dependency of PcIBMS1 and PcIPMS1 enzymatic activity on incubation conditions and components.** These enzymatic assays were performed using end-point enzyme assay (DTNB). These activities were measured with 0.5 mM 4-methyl-2-oxovalerate for PcIBMS1 and 0.5 mM 2-oxoisovalerate for PcIPMS1 in the presence of 0.5 mM acetyl-CoA and different concentration of divalent ions. To ensure optimum buffering capacity, 100 mM potassium phosphate buffer and 100 mM Tris-HCl buffer were used from pH 5-7 and pH 7.5-10, respectively. Divalent cations ($Ca^{2+}$, $Co^{2+}$, $Cu^{2+}$, $Mn^{2+}$, $Zn^{2+}$, $Mg^{2+}$) were used as chloride salts at a final concentration of 10 mM. To ensure optimum concentration of magnesium ion, the concentrations of magnesium ion ranging from 1mM to 100 mM were tested. The highest activity was set as 100%. The data are presented as relative mean percentages of three independent biological replicates. N.D. represents “not detected or relative activities < 5%”.

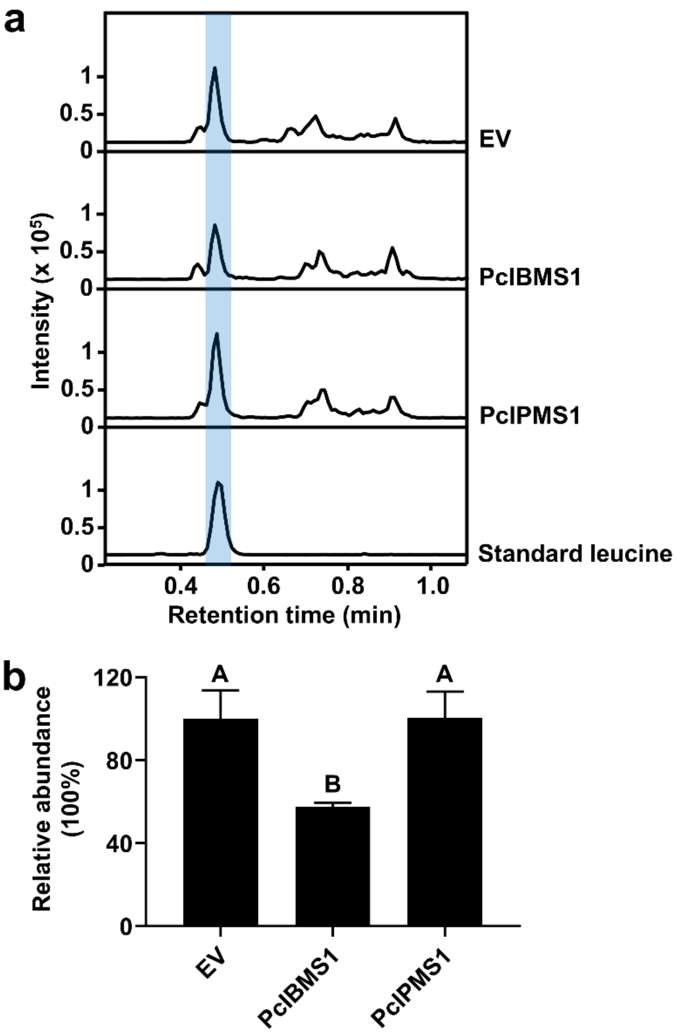


**Fig. S3 LC-QTOF-MS analyses of relative abundances of leucine in *N. benthamiana* leaves expressing *PcIBMS1* and *PcIPMS1*. a.** LC-QTOF-MS analyses of *N. benthamiana* leaves expressing the two enzymes PcIBMS1 and PcIPMS1. *N. benthamiana* leaves transformed with empty vector were used as negative control. Extracted ion chromatograms in negative mode of *m/z* 130.0874 for leucine are shown. Leucine was identified based on the comparison of retention time and mass spectrums with the authentic standard. **b.** The relative abundance of leucine in *N. benthamiana* leaves expressing *PcIBMS1* or *PcIPMS1* and *N. benthamiana* leaves transformed with empty vector. The relative abundance of leucine in transgenic *N. benthamiana* leaves was calculated by normalization of peak area of leucine to that of internal standard naringenin (*m/z*, 271.0612).

The relative abundance of leucine in *N. benthamiana* leaves transformed with empty vector was set as 100%. Significant difference among the relative abundance of leucine in transgenic *N. benthamiana* leaves was tested using independent *t*-tests $P < 0.01$ (Two-tailed distribution; Two-sample equal variance (homoscedastic)). Different letters above the bars mark the statistically significant groups after *t*-tests. EV, empty vector.

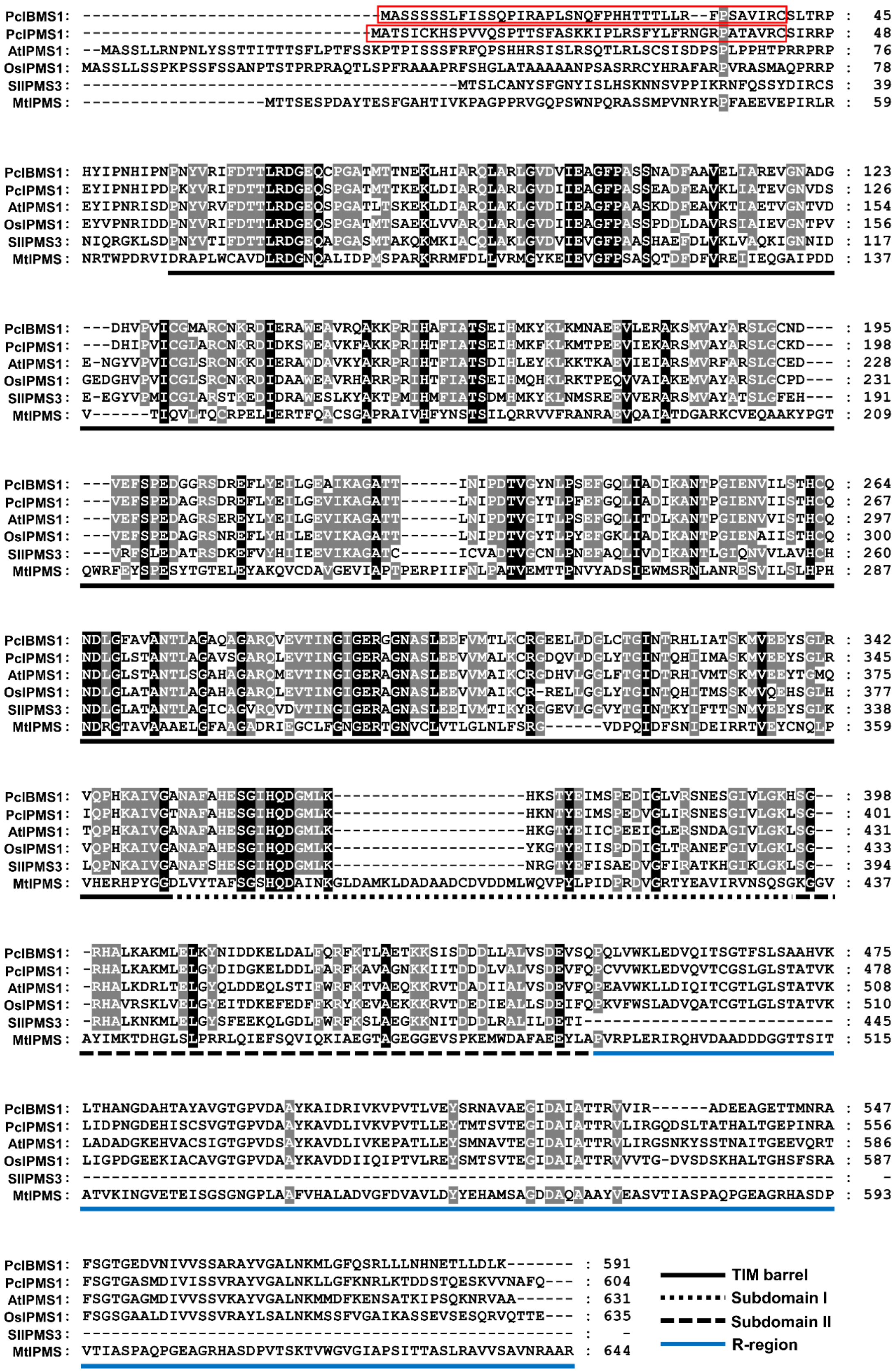
PcIBMS1: -------------------------------MASSSSSLFISSQPIRAPLSNQFPHHTTTLLR--FPSAVIRCSLTRP : 45
PcIPMS1: ------------------------------MATSICKHSPVVQSPTTSFASKKIPLRSFYLFRNGRPATAVRCSIRRP : 48
AtIPMS1: --MASSLLRNPNLYSSTTITTTSFLPTFSSKPTPISSSFRFQPSHHRSISLRSQTLRLSCSISDPSPLPPHTPRRPRP : 76
OsIPMS1: MASSLLSSPKPSSFSSANPTSTPRPRAQTLSPFRAAAPRFSHGLATAAAAANPSASRRCYHRAFARPVRASMAQPRRP : 78
SlIPMS3: ---------------------------------------MTSLCANYSFGNYISLHSKNNSVPPIKRNFQSSYDIRCS : 39
MtIPMS: -------------------MTTSESPDAYTESFGAHTIVKPAGPPRVGQPSWNPQRASSMPVNRYRPFAEEVEPIRLR : 59
PcIBMS1: HYIPNHIPNPNYVRIFDTTLRDGEQCPGATMTTNEKLHIARQLARLGVDVIEAGFPASSNADFAAVELIAREVGNADG : 123
PcIPMS1: EYIPNHIPDPKYVRIFDTTLRDGEQSPGATMTTKEKLDIARQLARLGVDIIEAGFPASSEADFEAVKLIATEVGNVDS : 126
AtIPMS1: EYIPNRISDPNYVRVFDTTLRDGEQSPGATLTSKEKLDIARQLAKLGVDIIEAGFPAASKDDFEAVKTIAETVGNTVD : 154
OsIPMS1: EYVPNRIDDPNYVRIFDTTLRDGEQSPGATMTSAEKLVVARQLARLGVDIIEAGFPASSPDDLDAVRSIAIEVGNTPV : 156
SlIPMS3: NIQRGKLSDPNYVTIFDTTLRDGEQAPGASMTAKQKMKIACQLAKLGVDVIEVGFPAASHAEFDLVKLVAQKIGNNID : 117
MtIPMS: NRTWPDRVIDRAPLWCAVDLRDGNQALIDPMSPARKRRMFDLLVRMGYKEIEVGFPSASQTDFDFVREIIEQGAIPDD : 137
PcIBMS1: ---DHVPVICGMARCNKRDIERAWEAVRQAKKPRIHAFIATSEIHMKYKLKMNAEEVLERAKSMVAYARSLGCND--- : 195
PcIPMS1: ---DHIPVICGLARCNKRDIDKSWEAVKFAKKPRIHTFIATSEIHMKFKLKMTPEEVIEKARSMVAYARSLGCKD--- : 198
AtIPMS1: E-NGYVPVICGLSRCNKKDIERAWDAVKYAKRPRIHTFIATSDIHLEYKLKKTKAEVIEIARSMVRFARSLGCED--- : 228
OsIPMS1: GEDGHVPVICGLSRCNKRDIDAAWEAVRHARRPRIHTFIATSEIHMQHKLRKTPEQVVAIAKEMVAYARSLGCPD--- : 231
SlIPMS3: E-EGYVPMICGLARSTKEDIDRAWESLKYAKTPMIHMFIATSDMHMKYKLNMSREEVVERARSMVAYATSLGFEH--- : 191
MtIPMS: V------TIQVLTQCRPELIERTFQACSGAPRAIVHFYNSTSILQRRVVFRANRAEVQAIATDGARKCVEQAAKYPGT : 209
PcIBMS1: ---VEFSPEDGGRSDREFLYEILGEAIKAGATT------INIPDTVGYNLPSEFGQLIADIKANTPGIENVILSTHCQ : 264
PcIPMS1: ---VEFSPEDAGRSDREFLYEILGEVIKAGATT------LNIPDTVGYTLPFEFGQLIADIKANTPGIENVIISTHCQ : 267
AtIPMS1: ---VEFSPEDAGRSEREYLYEILGEVIKAGATT------LNIPDTVGITLPSEFGQLITDLKANTPGIENVVISTHCQ : 297
OsIPMS1: ---VEFSPEDAGRSNREFLYHILEEVIKAGATT------LNIPDTVGYTLPYEFGKLIADIKANTPGIENAIISTHCQ : 300
SlIPMS3: ---VRFSLEDATRSDKEFVYHIIEEVIKAGATC------ICVADTVGCNLPNEFAQLIVDIKANTLGIQNVVLAVHCH : 260
MtIPMS: QWRFEYSPESYTGTELEYAKQVCDAVGEVIAPTPERPIIFNLPATVEMTTPNVYADSIEWMSRNLANRESVILSLHPH : 287
PcIBMS1: NDLGFAVANTLAGAQAGARQVEVTINGIGERGGNASLEEFVMTLKCRGEELLDGLCTGINTRHLIATSKMVEEYSGLR : 342
PcIPMS1: NDLGLSTANTLAGAVSGARQLEVTINGIGERAGNASLEEVVMALKCRGDQVLDGLYTGINTQHIIMASKMVEEYSGLR : 345
AtIPMS1: NDLGLSTANTLSGAHAGARQMEVTINGIGERAGNASLEEVVMAIKCRGDHVLGGLFTGIDTRHIVMTSKMVEEYTGMQ : 375
OsIPMS1: NDLGLATANTLAGAHAGARQLEVTINGIGERAGNASLEEVVMAIKCR-RELLGGLYTGINTQHITMSSKMVQEHSGLH : 377
SlIPMS3: NDLGLATANTLAGICAGVRQVDVTINGIGERAGNASLEEIVMTIKYRGGEVLGGVYTGINTKYIFTTSNMVEEYSGLK : 338
MtIPMS: NDRGTAVAAAELGFAAGADRIEGCLFGNGERTGNVCLVTLGLNLFSRG------VDPQIDFSNIDEIRRTVEYCNQLP : 359
PcIBMS1: VQPHKAIVGANAFAHESGIHQDGMLK--------------------HKSTYEIMSPEDIGLVRSNESGIVLGKHSG-- : 398
PcIPMS1: IQPHKAIVGTNAFAHESGIHQDGMLK--------------------HKNTYEIMSPEDVGLIRSNESGIVLGKLSG-- : 401
AtIPMS1: TQPHKAIVGANAFAHESGIHQDGMLK--------------------HKGTYEIICPEEIGLERSNDAGIVLGKLSG-- : 431
OsIPMS1: VQPHKAIVGANAFAHESGIHQDGMLK--------------------YKGTYEIISPDDIGLTRANEFGIVLGKLSG-- : 433
SlIPMS3: LQPNKAIVGANAFSHESGIHQDGMLK--------------------NRGTYEFISAEDVGFIRATKHGIKLGKLSG-- : 394
MtIPMS: VHERHPYGGDLVYTAFSGSHQDAINKGLDAMKLDADAADCDVDDMLWQVPYLPIDPRDVGRTYEAVIRVNSQSGKGGV : 437
PcIBMS1: -RHALKAKMLELKYNIDDKELDALFQRFKTLAETKKSISDDDLLALVSDEVSQPQLVWKLEDVQITSGTFSLSAAHVK : 475
PcIPMS1: -RHALKAKMLELGYDIDGKELDDLFARFKAVAGNKKIITDDDLVALVSDEVFQPCVVWKLEDVQVTCGSLGLSTATVK : 478
AtIPMS1: -RHALKDRLTELGYQLDDEQLSTIFWRFKTVAEQKKRVTDADIIALVSDEVFQPEAVWKLLDIQITCGTLGLSTATVK : 508
OsIPMS1: -RHAVRSKLVELGYEITDKEFEDFFKRYKEVAEKKKRVTDEDIEALLSDEIFQPKVFWSLADVQATCGTLGLSTATVK : 510
SlIPMS3: -RHALKNKMLELGYSFEEKQLGDLFWRFKSLAEGKKNITDDDLRALILDETI-------------------------- : 445
MtIPMS: AYIMKTDHGLSLPRRLQIEFSQVIQKIAEGTAGEGGEVSPKEMWDAFAEEYLAPVRPLERIRQHVDAADDDGGTTSIT : 515
PcIBMS1: LTHANGDAHTAYAVGTGPVDAAYKAIDRIVKVPVTLVEYSRNAVAEGIDAIATTRVVIR------ADEEAGETTMNRA : 547
PcIPMS1: LIDPNGDEHISCSVGTGPVDAAYKAVDLIVKVPVTLLEYTMTSVTEGIDAIATTRVLIRGQDSLTATHALTGEPINRA : 556
AtIPMS1: LADADGKEHVACSIGTGPVDSAYKAVDLIVKEPATLLEYSMNAVTEGIDAIATTRVLIRGSNKYSSTNAITGEEVQRT : 586
OsIPMS1: LIGPDGEEKIACAVGTGPVDAAYKAVDDIIQIPTVLREYSMTSVTEGIDAIATTRVVVTG-DVSDSKHALTGHSFSRA : 587
SlIPMS3: ------------------------------------------------------------------------------ : -
MtIPMS: ATVKINGVETEISGSGNGPLAAFVHALADVGFDVAVLDYYEHAMSAGDDAQAAAYVEASVTIASPAQPGEAGRHASDP : 593
PcIBMS1: FSGTGEDVNIVVSSARAYVGALNKMLGFQSRLLLNHNETLLDLK------- : 591
PcIPMS1: FSGTGASMDIVISSVRAYVGALNKLLGFKNRLKTDDSTQESKVVNAFQ--- : 604
AtIPMS1: FSGTGAGMDIVVSSVKAYVGALNKMMDFKENSATKIPSQKNRVAA------ : 631
OsIPMS1: FSGSGAALDIVVSSVRAYLSALNKMSSFVGAIKASSEVSESQRVQTTE--- : 635
SlIPMS3: --------------------------------------------------- : -
MtIPMS: VTIASPAQPGEAGRHASDPVTSKTVWGVGIAPSITTASLRAVVSAVNRAAR : 644
TIM barrel
Subdomain I
Subdomain II
R-region

**Fig. S4 Amino acid sequence alignment of PcIBMS1, PcIPMS1, AtIPMS1, OsIPMS1, SlIPMS3 and MtIPMS.** Black background shows perfectly conserved sequences across the IPMS proteins, while the light grey background shows the amino acid identity higher than 80% across the IPMS proteins. Red frames indicate the predicted chloroplast transit peptides of PcIBMS1 and PcIPMS1. Protein domain designations are derived from MtIPMS crystal structure (Koon *et al.*, 2004). PcIBMS1 and PcIPMS1 were directly obtained from *P. cablin* transcriptome database. The information of other proteins used in this sequence alignment is shown in Table S5.

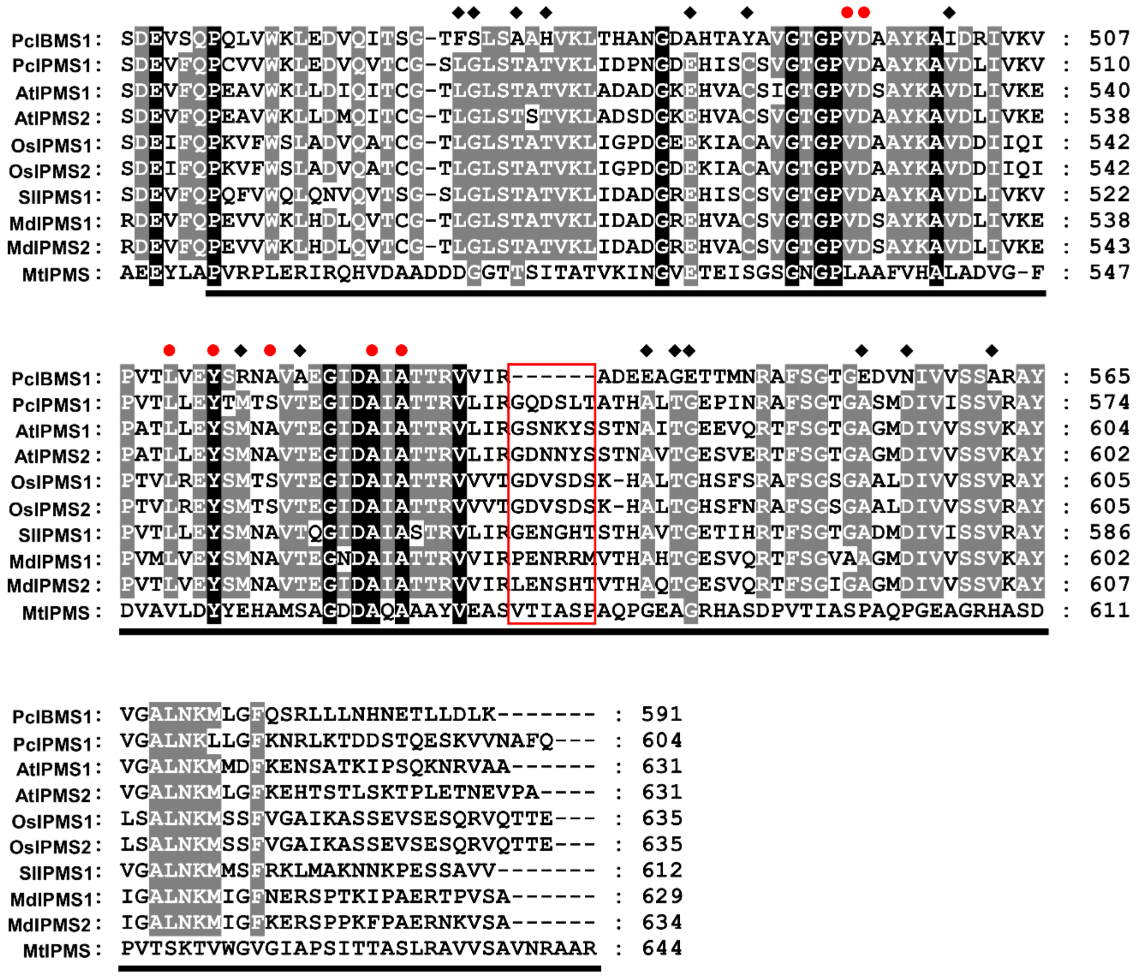


**Fig. S5 Sequence alignment of C-terminal domain of PcIBMS1, PcIPMS1, MtIPMS and other plant typical IPMSs that are subject to Leu feedback inhibition.** The IPMSs used in this sequence alignment except PcIBMS1 have been biochemically verified to be subject to Leu feedback inhibition (Koon *et al.*, 2004; de Carvalho *et al.*, 2005; de Kraker *et al.*, 2007; Ning *et al.*, 2015; He *et al.*, 2019; Sugimoto *et al.*, 2021). The sold diamond indicates amino acid divergence in the site of PcIBMS1 protein corresponding to the conserved amino acid site in the plant typical IPMSs occurs. Sold red circle marks the Leu binding sites that are derived from *Mycobacterium tuberculosis* IPMS (MtIPMS) crystal structure (Koon *et al.*, 2004). Red frame indicates the sites where PcIBMS1 loss five amino acids. Blank lines under the sequence alignment depict the C-terminal allosteric regulatory domain based on the MtIPMS crystal structure (Koon *et al.*, 2004). Black background shows perfectly conserved sequences across the IPMS proteins, while the light gray background shows the amino acid identity higher than 80% across the IPMS proteins. PcIBMS1 and PcIPMS1 were directly obtained from *P. cablin* transcriptome database. The information of other proteins used in this sequence alignment is shown in Table S5.
